# Essential genes are dominantly activated by single transcription factors

**DOI:** 10.64898/2026.04.27.721063

**Authors:** Martina Capriati, Sylke Nestmann, Izlem Su Akan, Faustus Benedikt Tuschmann, Silke Druffel-Augustin, Ralph S. Grand

## Abstract

Cell viability depends on the precise expression of essential genes, which are controlled by CpG-island (CGI) promoters densely bound by transcription factors (TFs). This has led to the prevailing model that TFs cooperate to ensure ubiquitous expression. Here, using rapid and reversible single and combinatorial degradation in murine stem cells, we systematically dissect the regulatory interactions between five key TFs. We uncover an unexpectedly specific architecture in which regulatory dominance, rather than cooperation, is the prevailing mode, where individual TFs autonomously drive chromatin opening and gene activation at largely distinct promoters. Cooperative regulation occurs at a minority of sites with antagonistic or synergistic outcomes modulated by the interplay between nucleosome positioning and TF sensitivity to chromatin. This logic is recapitulated at synthetic sequences and reflected in human genetic variation. These findings reveal that single TFs dominantly activate distinct sets of CGI-linked genes, including essential genes, across development, homeostasis, and disease.

## Introduction

Cellular survival depends on the expression of genes that encode for fundamental biological processes, such as metabolism and the cell cycle. These common essential genes (CEGs) are ubiquitously expressed, broadly conserved across species, and display reduced coding-sequence variation^1,2^. Although ubiquitously required by all cell types, their misregulation is associated with disease, for example, the recurrent overexpression of cMYC in cancer or the diminished activity of metabolic pathways during aging^3–5^. Despite their importance, the essential nature of these genes has limited the efforts to dissect the mechanisms governing their transcriptional regulation. CEGs are predominantly driven by CpG island (CGI) promoters, which are the major promoter class in human cells and account for over 70% of RNA polymerase II-dependent transcripts^6,7^. CGIs are distinct in that they frequently display open chromatin and active histone modifications (H3K4me3, H3K27ac) across cell types^7^. This pervasive chromatin state has fostered the prevailing view that CGIs are intrinsically permissive elements that ensure ubiquitous transcription by enabling multiple TFs to cooperative or act redundently to regulate gene activity.

TFs are central to gene regulation, recognizing short DNA motifs (6-25 bp) within cis-regulatory elements (CREs) to control chromatin state and transcription^8,9^. Although mammalian genomes contain millions of potential TF motifs, only a small subset is occupied in any given cell type^10^. Chromatin organization is a key determinant of this selectivity: most TFs are chromatin-sensitive and cannot efficiently engage nucleosomal DNA alone^11,12^. By contrast, it is thought that a limited class of chromatin-insensitive TFs (pioneer factors) can bind nucleosomal DNA directly, remodel local chromatin, and recruit additional factors^13,14^. The combinatorial interplay among TFs with different sensitivities to chromatin is thought to overcome the nucleosome barrier and shape CREs activity^15^. This cooperativity is facilitated by the precise arrangment of motifs enabling TFs to achieve enhanced DNA-binding and stimulate the opening of chromatin and gene activiation at shared cis-regulatory elements^16–19^Canonical examples include OCT4-SOX2, which engages a composite motif critical for pluripotency^20–22^ and the more recently described TWIST1-ALX4 pair, which binds the coordinator motif to orchestrate mesenchymal gene expression^17^. Alternatively, redundant TF binding provides robustness in transcriptional control, buffering gene expression against alterations in DNA sequence or TFs. Houskeeping gene promoters, which often contain multiple overlapping motifs and TF binding events, are thought to be controled by both cooperative and redundant TF activity. However, a detailed mechanistic understanding of how TFs cooperate in chromatin to regulate housekeeping gene expression is lacking, particularly at CEG promoters.

Here, we systematically dissect the regulatory logic controlling CEG expression by focusing on critical TFs that co-occupy their promoters. To overcome the lethality of removing essential regulators, we used inducible degron systems to achieve rapid depletion and recovery of individual as well as combinations of TFs. By integrating immediate (within 1 hour) changes in chromatin accessibility, TF binding, and transcription following depletion, we uncover gene subsets that are dominantly regulated by single TFs despite extensive co-binding. Our analyses revealed limited cooperative or redundant activity instead there is a high level of specificity at CGI promoters, where a single dominant TF governs promoter accessibility and transcriptional output. Co-bounding of TFs is modulated by chromatin around the dominant TF, occationally tuning gene expression, potentially enabling the integration of cell type specific signals. We confirm this single-factor dominance and chromatin mediated gene activity modulation with synthetic constructs and deep learning models indicate conservation in human cells. These findings redefine our understanding of CGI regulation, revealing that cell viability is governed by dominant TFs and enabling to link disease-related noncoding variation to effectors.

## Result

### Tight control of CEG expression levels distinguishes cell identity

Different cell types exhibit distinct functional and metabolic demands^23^. While these differences are typically attributed to tissue-specific gene expression, we hypothesized that the precise expression level of CEGs also contributes to defining cell identity. If true, CEG expression levels should be sufficient to distinguish cell types, even within complex tissues such as the brain. To test this, we first compiled a high-confidence set of approximately 1,800 CEGs by integrating gene essentiality datasets from healthy and cancer cell types^1,24,25^. We then analyzed their expression levels in single-cell RNA-sequencing data from Tabula Muris, and found substantial variation across cell types (Extended data Fig.1a). Remarkably, the different level of CEGs alone was sufficient to distinguish individual cell types in the brain with an accuracy comparable to that achieved using the entire transcriptome (Figure 1a, Extended data Fig.1b). This finding indicates that, despite their ubiquitous requierment, CEG expression level is cell type-specific and must be tightly controlled, motivating a mechanistic dissection of how they are regulated.

**Figure 1.**
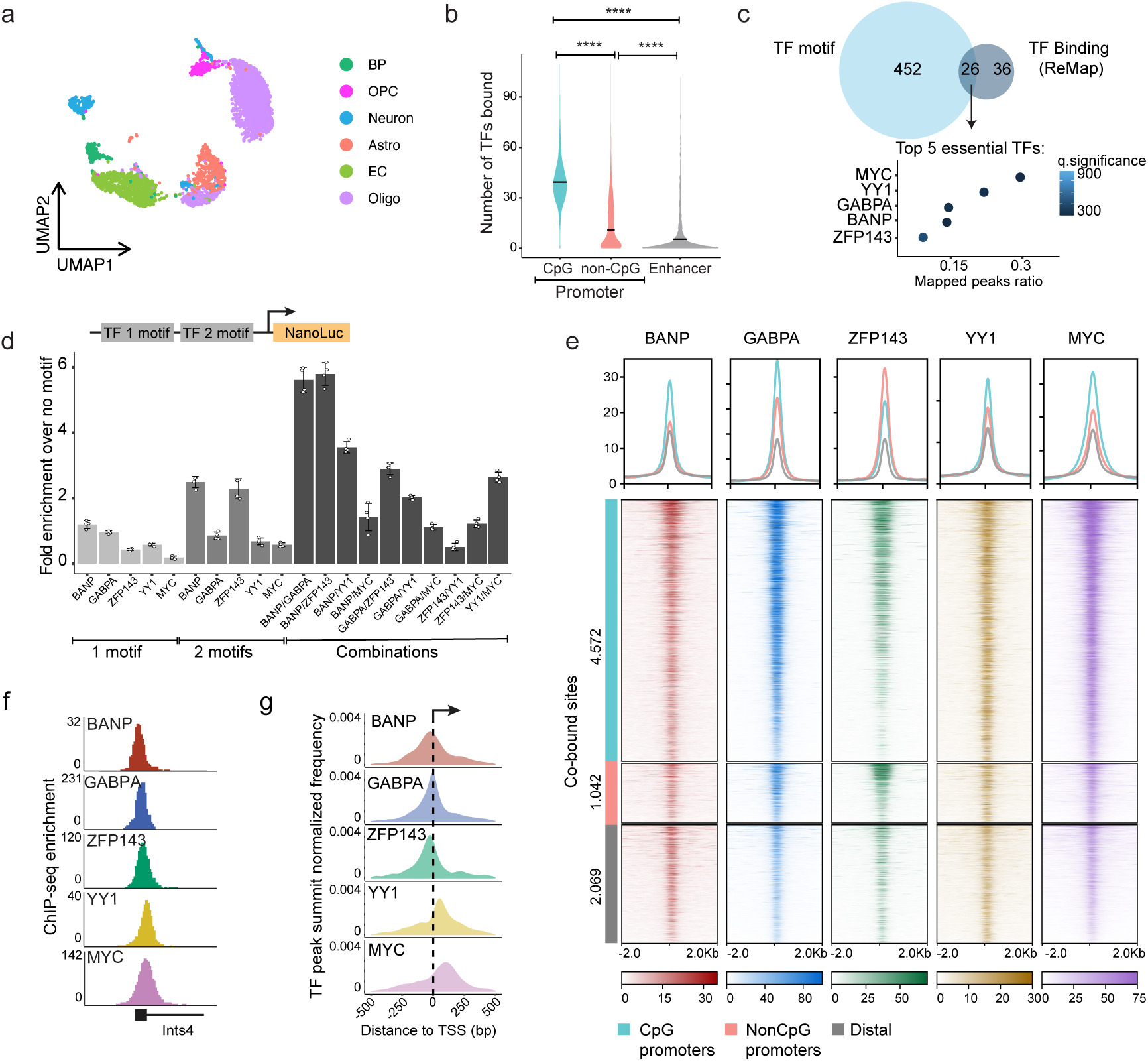
TFs that regulate CEGs co-bind extensively around the TSS at CGI promoters. a) Differential expression of CEGs in a complex tissue like the brain can distinguish cell types. The Tabula Muris single-cell RNA-sequencing data was used for the analysis. b) Number of TFs bound at distinct cis-regulatory elements. Data obtained from ReMap. c) The fraction of TF binding that overlaps with CEG promoters using ReMap data, and also have a motif; the top 5 essential TFs are displayed. d) Activity of a Nano-Luciferase gene driven by synthetic promoters harboring different combinations of the consensus motifs of the five TFs we selected for detailed investigation. e) Binding enrichment of the five TFs at co-bound sites, divided into different cis-regulatory elements, displayed as a waterfall plot (bottom) with a cumulative metaplot (top). f) Single locus example of ChIP-seq enrichment at a CEG that is co-bound by all five of the TFs. g) Positional binding preference of the five TFs at co-bound CGI promoters relative to the TSS. Density plot: for each category, we generated a smoothed distribution of binding positions by computing a univariate KDE of distances from the TSS (−250 to +250 bp) with 200 evaluation points. Vertical reference lines were drawn at the TSS (0 bp; solid) and at ±30 bp (dotted). *** = pvalue<0.0001.

The majority of CEGs are controlled by CGI promoters that have the largest number and diversity of TFs bound compared to non-CGI promoters or enhancers (Figure 1b, Extended data Fig.1c,d). This suggests a complex regulatory network where TF cooperation and redundancy likely underlie the ubiquitous and precisely tuned expression of essential genes.

### Extensive transcription factor co-binding characterizes CGI promoters

To interrogate the regulatory logic at CGI promoters, with a focus on CEGs, we sought to nominate TFs for systematic dissection based on motif presence and binding. We performed TF motif enrichment analysis and intersected this with 81 publicly available chromatin immunoprecipitation followed by sequencing (ChIP-seq) datasets in mouse embryonic stem cells (mESCs) to rank binding frequency at CEG promoters^26,27^. We chose to focus on the top five ranked essential TFs, yielding: BANP, GABPA, ZFP143, YY1, and MYC, which represent a range of DNA binding domain classes and chromatin sensitivities (Figure 1c, Extended data Fig.1e)^8,12^. The general relevance of these TFs is highlighted by work showing their strong association with open chromatin across cell types during human development^28–30^, and that motif mutations result in modified gene expression, with relevance for disease^31^.

To assess their regulatory potential, we constructed synthetic promoters containing single or pairwise combinations of consensus binding motifs for the selected TFs in a CGI-like sequence context^12,27,32^. These were cloned upstream of a Nano-Luciferase (NanoLuc) reporter gene and transiently transfected into mESCs. Single motifs mildly increased reporter activity, with BANP showing the strongest effect, whereas homo- or heterologous pairs of TF motifs synergistically enhanced transcription, particularly those involving BANP (Figure 1d). These results support that TFs cooperate to activate CGI promoters, with combinations of multiple TFs amplifying promoter activity.

To dissect the regulatory network of these TF, we generated homozygous degron (see below) together with V5-affinity tag knock-in mESC lines for each TF using CRISPR/Cas9 (Figure 2a, Extended data Fig.2a)^33^. To ensure that the addition of these tags has not disrupted their genomic occupancy and define a set of promoters where they co-bind, we sought to map their localization in our system. We performed ChIP-seq and identified high-confidence consensus peaks for each factor from highly reproducible (Pearson correlation >=0.93) replicate experiments (BANP, 3,659; GABPA, 11,138; ZFP143, 7,165; MYC, 10,196; YY1, 8,577), with strong enrichment for the respective TF motif and conserved genomic distribution (Extended data Fig.1f,g)^27,34–37^. For statistical robustness and comparability for subsequent analysis, we merged peaks within 200 bp, yielding a set of 29,288 regions, roughly split between promoters (mostly CGIs) and distal sites (Extended data Fig.1h,i). This identified extensive co-binding at CGI promoters (∼50%), including at a large fraction of CEGs (Figure 1e,f, Extended data Fig.1j-l). Binding was clustered tightly around the transcription start site (TSS), with distinct positional preferences; BANP, GABPA, and ZFP143 bound predominantly upstream of the TSS, whereas YY1 and MYC were enriched downstream (Figure 1g)^38–40^.

**Figure 2.**
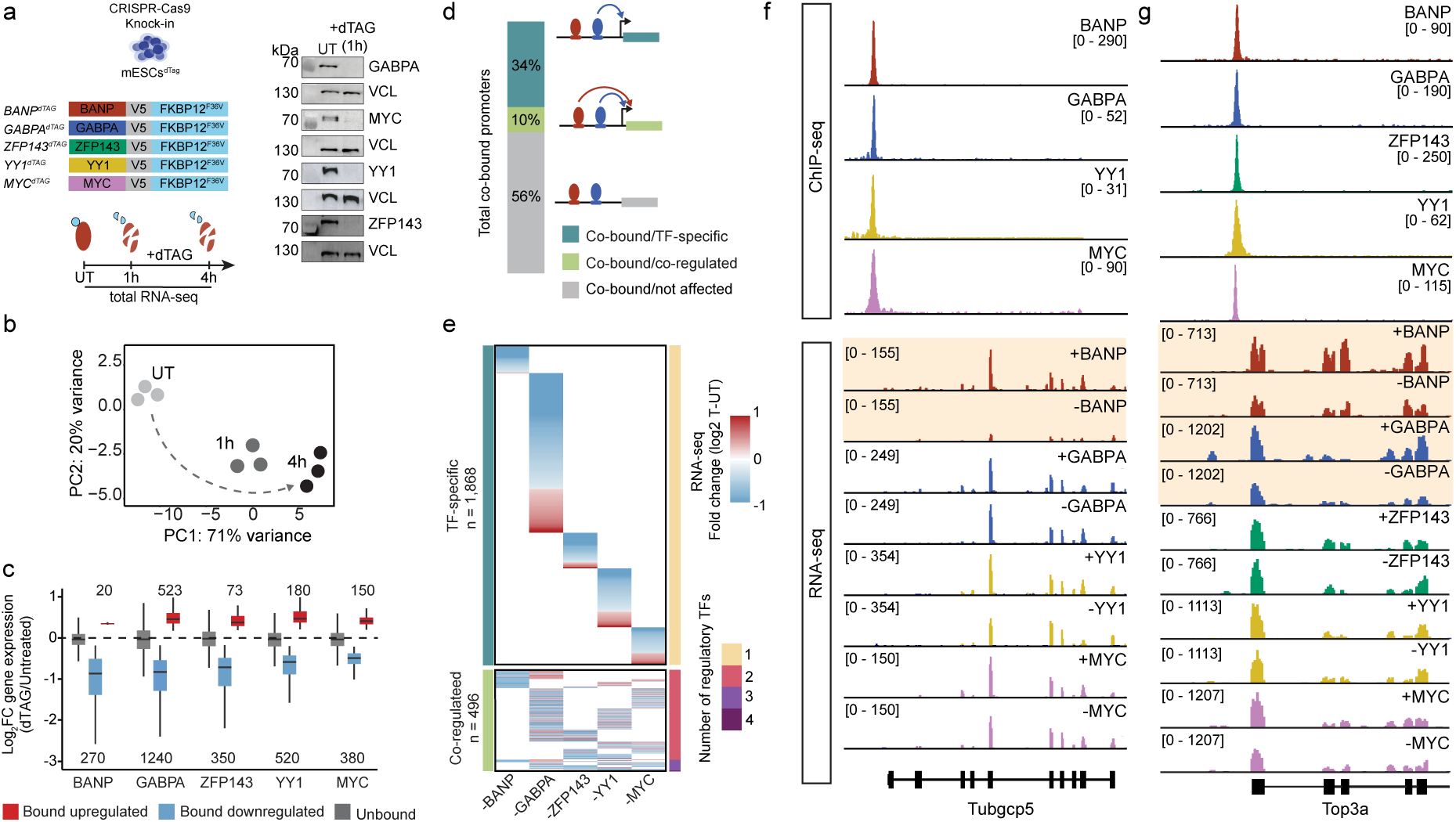
Co-bound genes are predominantly controlled by a single TF. a) Schematic of the dTAG knockin strategy and Western blot of homozygous knockin clones showing depletion following one hour of degradation upon dTAG13 treatment. RNA-seq was performed 1h and 4h post-degradation. b) Principal Component Analysis (PCA) of RNA-seq data showing progressive transcriptional divergence over time after TF degradation relative to untreated (UT). c) The number and log-fold change in transcription of genes following the acute depletion of each TF. d) Schematic of our classification of co-bound genes into different RNA response groups and the fraction of co-bound genes that fall into the different response groups shown in. e) The log-fold change in gene transcription at co-bound genes that responded to the acute depletion of only one of the TFs or more is displayed as a heatmap. Genes with a log-fold change greater than +/- 1 were capped. f) Single locus example of a CEG that is co-bound by four of the TFs, but where gene activity is only affected by the removal of one of the TFs. Genome browser tracks of ChIP-seq (top) and RNA-seq (bottom) are displayed. g) Single locus example of a CEG that is co-bound by five of the TFs, and where gene activity is affected by the removal of two of the TFs. Genome browser tracks of ChIP-seq (top) and RNA-seq (bottom) are displayed.

These findings show that CGI promoters, including CEGs, are densely co-bound by TFs in proximity to the TSS, suggesting a highly integrated regulatory architecture. Together with their cooperative potential in synthetic promoters, this indicates that these factors likely act together to regulate transcription.

### Co-bound genes are predominantly activated by individual TFs

To determine the impact of individual TFs on transcription and whether co-bound promoters are also co-regulated, we sought to capture the direct, primary response following their removal. Given that these TFs are all classified as essential, conventional knockout approaches were not feasible. Thus, we employed an inducible degron system that allowed acute TF depletion, enabling us to capture direct transcriptional effects^41^. We generated homozygous degron-tagged cell lines together with the V5-tag (see above, Figure 2a), enbaling to induce the rapid depletion of each TF within one hour of addition of the PROTAC compound (Figure 2a, Extended data Fig.2a). Prolonged depletion led to reduced cell viability or death for all TFs, consistent with their essential roles in cellular maintenance (Extended data Fig.2b)^27,35^.

To capture primary transcriptional responses, we performed total RNA-seq after 1- and 4-hours of TF depletion, and determined significant changes in nascent transcript levels using intron-exon split analysis (see Methods, Figure 2b, Extended data Fig.2c,d)^42^. The majority of differentially expressed genes were strongly bound by the corresponding TF, confirming that they were direct targets (Extended data Fig.2d,e), yet only a fraction (10-30%) of bound promoters exhibited transcriptional changes (Extended data Fig.2f), as seen in other systems/organisms^43,44^. All TFs function predominantly as transcriptional activators, with strong downregulation of their target genes following depletion (Figure 2c, Extended data Fig.2d). In line with their central role in cell viabiltiy, these five TFs collectively regulate the activity of 34% of CEGs, including key components of the cell cycle, ribosome biogenesis, and mitochondrial biogenesis (Extended data Fig.2g,h).

We next examined whether co-binding of multiple TFs resulted in the co-regulation of the same gene. Focusing on the ∼4,500 co-bound CGI promoters (Figure 1e,2d), we found that transcriptional responses were surprisingly dominated by single TFs. Approximately 1,800 genes (34% of co-bound genes) were regulated by one TF, whereas only about 10% (∼500 genes) responded to the depletion of more than one TF (Figure 2e-g, Extended data Fig.2k), indicating that co-binding infrequently translates into co-regulation of transcription. Moreover, the magnitude of downregulation was comparable or greater for genes that responded to the removal of one TF compared to co-regulated genes (Extended data Fig.2i), suggesting that a single TF often exerts the major regulatory influence at co-bound promoters. This could not be explained by the difference in binding strength, with co-regulated genes exhibiting similar or even higher TF occupancy compared to TF-specific regulated genes (Extended data Fig.2j). Sorting co-regulated genes by the number of TFs that influence their expression showed that most were controlled by combinations of two TFs, despite additional co-binding (Figure 2e,g, Extended data Fig.2k). Interestingly, these strong activating TFs can have both synergistic and antagonistic effects on transcription at co-regulated genes, indicating locus-specific interactions that may tune transcriptional output. These results reveal that individual TFs can activate transcription from co-bound promoters, and raise the question of why multiple essential TFs co-occupy the same CGI promoters.

### TFs control chromatin state at distinct CGI promoters

We therefore next asked whether TF co-occupancy might reflect distinct contributions to promoter chromatin architecture rather than direct co-regulation of transcription. We examined whether the TFs shape chromatin states by profiling chromatin accessibility and histone modifications at matched time points (within 1 hour) following acute TF depletion (Figure 3a). To measure changes in open chromatin, we performed Assay for Transposase-Accessible Chromatin using sequencing (ATAC-seq) following 1- and 4-hours of TF depletion (Extended data Fig.3a)^45^. Removal of BANP, GABPA, or ZFP143 resulted in rapid changes in chromatin accessibility at a subset (5-10%) of their most strongly bound sites, with most effects occurring within the first hour (Figure 3b, Extended data Fig.3b, c, g). These changes occurred at distinct sites, with a strong loss of accessibility at CGI promoters, despite extensive co-binding (Figure 3c,d, Extended data Fig.3d-f). These results suggest that even within shared binding landscapes, individual TFs maintain chromatin accessibility at distinct promoters. Some sites also showed increased accessibility, mainly at distal regulatory elements (Extended data Fig.3d,e). By contrast, YY1 and MYC depletion caused minimal changes in chromatin accessibility, despite producing strong transcriptional effects (Figure 3b).

**Figure 3.**
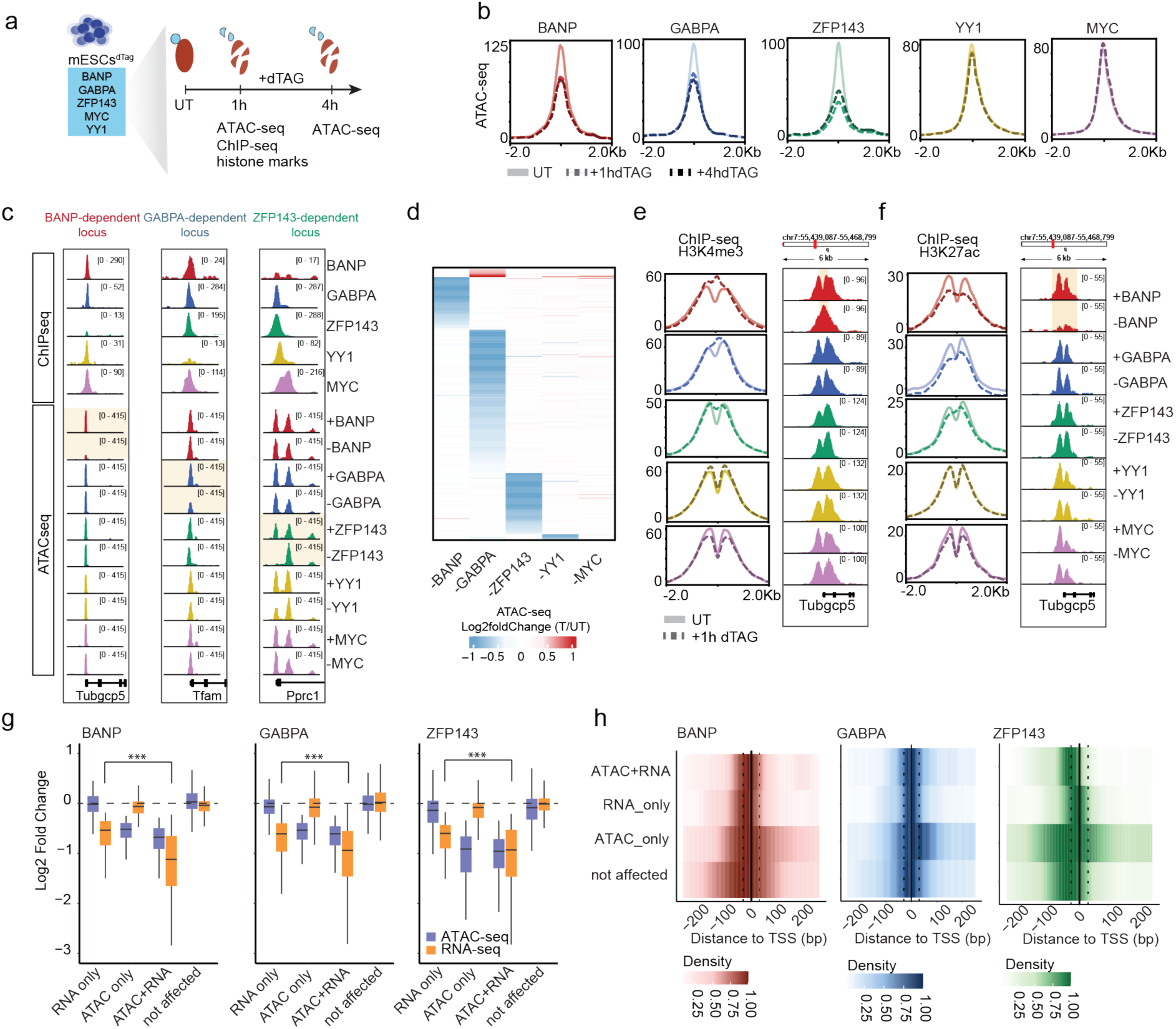
TF specific chromatin organization and modification. a) Schematic of how we captured the primary changes in chromatin organization and modification to the removal of individual TFs. b) Changes in chromatin accessibility following the acute depletion of each TF. The metaplots show the change in accessibility at bound and significantly changing sites for BANP, GABPA, and ZFP143, and just bound sites for YY1 and MYC, which do not significantly impact open chromatin. c) Single locus examples of co-bound CEG promoters where only one of the TFs impacts chromatin accessibility. d) The heatmap shows the fold change (log2) in open chromatin at all the sites where there were significant changes in chromatin accessibility in all the cell lines. e and f) Metaplots showing the change in enrichment and distribution of epigenetic marks on the same regions displayed for the chromatin accessibility plots (Left). Representative locus showing histone marks ChIP-seq signal at Tubgc5 promoter, a BANP-regulated site (Right). g) The level of gene expression or chromatin accessibility change in the different classes of genes. h) Binding position of each TF relative to the TSS, divided into the different gene classes as in (g). *** = p<0.0001.

This observation for YY1 and MYC prompted us to investigate whether additional chromatin features, such as active histone marks, better reflect TF transcriptional activity. We profiled Histone 3 Lysine 4 trimethylation (H3K4me3) and Histone 3 Lysine 27 acetylation (H3K27ac) before and after one hour of TF depletion. Loss of BANP, GABPA, or ZFP143 produced rapid and localized changes in both marks (Figure 3e,f). H3K4me3 level remained globally stable, but the enrichment at the -1 and +1 nucleosomes was redistributed over the TF-bound site, consistent with nucleosome invasion following TF loss (Figure 3e)^27^. By contrast, H3K27ac levels were markedly reduced, most prominently following BANP depletion (Figure 3f, Extended data Fig.3h), mirroring the loss of promoter activity. YY1 and MYC depletion, on the other hand, did not alter the level or distribution of either histone mark at promoters that showed a transcriptional effect (Figure 3e,f, Extended data Fig.3h), indicating that changes in histone modifications track more closely with a TF’s ability to remodel chromatin than regulate transcription.

This identified that only a subset of TFs: BANP, GABPA, and ZFP143, lead to the loss in open chromatin and altered histone mark landscapes at distinct CGI promoters. YY1 and MYC, by contrast, primarily function as transcriptional activators that likely require pre-established open chromatin. Together with the transcriptional response, this points to a hierarchical division of labor among co-bound TFs, where some can open chromatin and activate genes while others only modulate transcription within pre-existing active chromatin landscapes.

### TF location relative to the TSS links open chromatin to transcription

To explore the link between open chromatin and transcriptional regulation, we segregated the effected promoters following BANP, GABPA, or ZFP143 depletion into three categories: (1) those affected only at the transcriptional level (RNA-only), (2) only at the chromatin level (ATAC-only), and (3) those showing coordinated changes in both accessibility and transcription (ATAC+RNA) (Figure 3g, Extended data Fig.3i). Across all factors, the largest category comprised RNA-only responses (∼75%), while the remaining sites were roughly divided between ATAC-only and ATAC+RNA affected promoters (Extended data Fig.3j). Notably, genes in the ATAC+RNA group exhibited significantly stronger transcriptional changes compared to the RNA-only targets (Figure 3g), indicating that chromatin remodeling and transcriptional impact are coupled at the strongest responsive genes.

To uncover what distinguishes these regulatory outcomes, we analyzed both TF motif variants and binding position relative to the TSS. We performed motif scanning across all bound sites, derived distinct position weight matrices (PWMs) for each TF at affected (ATAC+RNA, RNA-only, and ATAC-only) compared to non-affected sites, and quantified differences in motif scores (see Methods). For BANP, there was a clear distinction with the optimal motif variant preferentially enriched at affected sites, most strongly in the ATAC+RNA group (Extended data Fig.3k). By contrast, there was little difference between the motif variant found at affected versus non-affected sites for GABPA and ZFP143. We then wondered if the binding position relative to the TSS might distinguish between the different functional classes. Indeed, BANP, GABPA, and ZFP143 binding at ATAC+RNA and RNA-only regulated promoters was sharply concentrated immediately upstream of the TSS, whereas ATAC-only or nonresponsive promoters showed more dispersed binding (Figure 3h). Furthermore, the effect of each TF on gene expression was strongest when the respective factor was positioned closest to the TSS relative to its co-bound partners (Extended data Fig.3l). Notably, this positional dependence was observed only for the chromatin-opening factors and not for YY1 or MYC, whose impact on RNA levels appeared largely independent of their position relative to the TSS (Extended data Fig.3l), suggesting that this effect is specifically linked to chromatin-opening activity. This positional bias is in line with previous findings that TF occupancy near the TSS is critical for transcriptional activation, with the BANP motif enriched ∼30 bp upstream of the TSS in reporter assays^39^, and both BANP and ZFP143 enhancing expression when bound in this region^38^. Interestingly, we also identified promoters where TF depletion altered chromatin accessibility but not gene expression. These sites were typically positioned farther from the TSS, suggesting that open chromatin alone is insufficient for productive transcription when misaligned with the core promoter architecture.

Together, our results suggest that a single TF can be responsible for both open chromatin and transcription at distinct subsets of CGI promoters, even when extensively co-binding by equally potent TFs. However, single-factor perturbations cannot account for functional redundancy or context-dependent cooperation among TFs.

### Open chromatin at co-bound promoters is primarily maintained by single TFs

To directly test for redundant and cooperative TF interactions requires the ability to independently and simultaneously deplete TF pairs, potentially using a double-degron system. We sought a degron tag that was comparable in size and displayed matched degradation and recovery kinetics to the dTAG, and chose the recently developed BromoTag degron39 40. To test the double-degron system, we focused on two dominant TFs and introduced the Bromo- and HA-tag into the *Banp* locus in the existing GABPA dTAG cell line using CRISPR/Cas9 (Figure 4a). This enabled the synchronized depletion and recovery of individual as well as both TFs (Figure 4b). To more generally explore TF cooperation, we generated additional double-degron cell lines, centered around BANP, which showed the strongest cooperative potential in reporter assays (Figure 1d). This resulted in four double-degron cell lines (BANP/GABPA, BANP/ZFP143, BANP/YY1, and BANP/MYC), in which individual or pairs of TFs can be removed and recovered synchronously (Figure 4b, Extended data Fig.4a).

**Figure 4.**
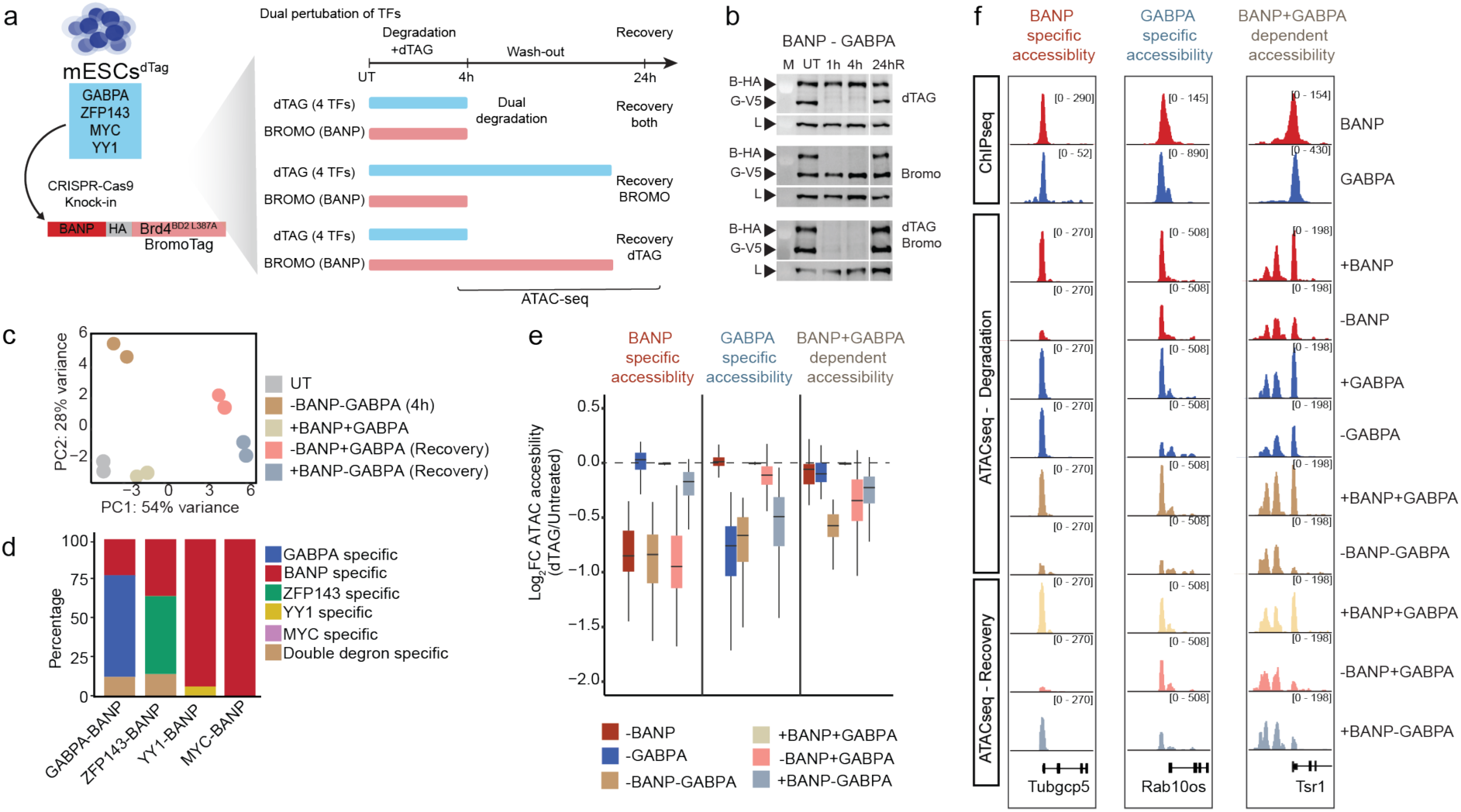
Generating open chromatin is driven by individual TFs. a) Schematic overview of the experimental workflow. mESCsdTAG expressing Bromo-tagged BANP were generated using CRISPR/Cas9-mediated genome editing. Individual or combinatorial degradation was induced using dTAG13 and/or AGB1 for 1h or 4h, followed by combinations of recovery timepoints. ATAC-seq and RNA-seq were performed to assess chromatin accessibility and gene expression dynamics. b) Western blot analysis of BANP and GABPA protein levels after 1 h, 2 h, 4 h of dTAG13 and/or AGB1 treatment, demonstrating efficient protein depletion and recovery. Vinculin (VCL) was used as a loading control. c) PCA of ATAC-seq data showing accessibility divergence after both TF degradation, recovery of only GABPA, recovery of only BANP, and recovery of both TFS, relative to untreated (UT). d) Stacked barplot showing the relative proportion of regulatory regions classified into BANP-specific, GABPA-specific, BANP+GABPA-dependent, results are also indicated for BANP-ZFP143 double degron experiment. e) IGV tracks showing representative loci (Tubbcp5, Rab10os, Tsr1) classified into three categories based on transcription factor dependence: BANP-specific, GABPA-specific, and BANP+GABPA-dependent accessibility. ChIP-seq profiles for BANP and GABPA are shown alongside ATAC-seq signal across untreated, degraded, and recovered conditions. f) Boxplots displaying log₂ fold-changes in accessibility (ATAC-seq) for peaks classified as BANP-specific, GABPA-specific, or BANP+GABPA-dependent following individual and combinatorial degradation and recovery conditions. Centre line, median; box limits, upper and lower quartiles; whiskers, minimum and maximum.

First, we tested whether TF pairs cooperate to maintain open chromatin by measuring whether combinatorial degradation resulted in the loss of chromatin accessibility at additional co-bound sites. The removal of BANP together with GABPA or ZFP143 uncovered a small set (∼10%) of new regions showing loss of chromatin accessibility that were non-significantly affected by the depletion of single factors (Figure 4c,d, Extended data Fig.4b,c). These represent loci where cooperative activity between chromatin-remodeling TFs is required to maintain open chromatin. However, the magnitude of accessibility loss upon dual depletion was modest, suggesting that true redundancy among chromatin-opening TFs is rare (Figure 4e,f, Extended data Fig.4c). The co-depletion of BANP together with YY1 or MYC did not result in additional effects, consistent with their lack of chromatin remodeling activity (Figure 3b,4d, Extended data Fig.4d,e). These results demonstrate that while cooperative maintenance of accessibility is observed at a few sites, open chromatin at the majority of co-bound promoters is primarily maintained by single TFs.

### Open chromatin is predominantly established by single TFs

The dominant maintenance of open chromatin by individual TFs, even at co-bound promoters, does not exclude a co-dependence to establish chromatin accessibility. To dissect the regulatory logic responsible for generating open chromatin at co-bound promoters, we next investigated the interdependence of TFs during chromatin re-engagement. The degron systems enabled us to directly test this by performing combinatorial TF degradation and recovery. For this, we optimized conditions to restore proteins to wild-type levels after removal of the degradation compound(s). Specifically, we depleted both TFs for four hours and then allowed individual or simultaneous recovery for 24 hours (24hR; Figure 4a,b, Extended data Fig.4a,b). Reintroduction of both TFs fully restored chromatin accessibility to levels comparable to undegraded cells, including at cooperative sites identified in the double-degron experiments (Figure 4c,e,f, Extended data Fig.4c,d,e). Interestingly, at sites dominated by BANP, GABPA, or ZFP143, reintroduction of the corresponding TF also re-opened chromatin, regardless of whether the partner factor was also restored. Remarkably, this demonstrated that each TF can autonomously re-establish accessibility at its target loci and cannot be compensated by the presence of an equally capable factor, consistent with context-dependent pioneer-like activity (Figure 4e,f, Extended data Fig.4c). By contrast, cooperative loci, where dual depletion results in accessibility loss, could only be partially re-opened when either TF was restored alone. Full recovery required re-expression of both TFs, indicating that these sites depend on combinatorial action for the establishment of open chromatin, which can then be redundantly maintained (Figure 4e,f, Extended data Fig.4c). Interestingly, BANP reintroduction also partially restored accessibility at certain GABPA- or ZFP143-dependent loci, whereas the reverse was not true (Figure 4e, Extended data Fig.4c). This asymmetry highlights BANP’s dominant position within the chromatin-opening hierarchy, capable of independently initiating and maintaining accessibility(Mol Cell Grand REF).

We further tested whether BANP’s ability to re-establish accessibility extends beyond chromatin-remodeling partners by examining its dependency on YY1 or MYC, two TFs that lack chromatin-opening activity (Figure 3b). Consistently, BANP reintroduction alone was sufficient to restore accessibility at BANP-regulated sites even in the continued absence of YY1 or MYC, confirming that these factors do not contribute to chromatin establishment (Extended data Fig.4d,e). This reinforces the conclusion that YY1 and MYC function as transcriptional amplifiers acting on pre-opened chromatin. These experiments reveal that the establishment of open chromatin at co-bound CGI promoters is often driven by individual TFs acting autonomously at distinct target sites. Thus, both the *de novo* formation and maintenance of open chromatin at essential gene promoters depend primarily on the activity of one dominant TF per locus. While at most promoters the TFs act independently, there is a subset that requires combinatorial input or relies on the prior action of a more potent TF. This potentially implies that many co-binding events depend on a dominant chromatin opener to expose binding sites.

### Pioneer activity establishes a binding hierarchy at co-bound promoters

Given that most co-binding events did not contribute to chromatin organization or gene transcription, we next asked whether the binding of additional TFs depends on chromatin-opening factors. Specifically, we tested whether YY1 and MYC, which lack remodeling activity, require prior open chromatin generated by pioneer TFs (BANP, GABPA, or ZFP143) to bind co-occupied promoters. To assess this, we first analyzed our ATAC-seq data for BANP, GABPA, and ZFP143 using TOBIAS to quantify changes in motif occupancy upon TF degradation^47^. This confirmed that the largest footprint reduction occurred at motifs for the degraded TF and also identified additional motif classes that showed reduced occupancy (Figure 5a, Extended data Fig.5a,b), indicative of a binding dependency. To directly measure the immediate effects on co-binding, we utilized our degron lines to perform ChIP-seq for the TFs before and after acute (1 h) depletion of each chromatin-opening TF (Figure 5b, Extended data Fig.5c). At sites where BANP, GABPA, or ZFP143 drive open chromatin, their removal led to a pronounced decrease in YY1 and MYC binding at co-bound CGI promoters (Figure 5c,d), indicating that these chromatin-sensitive TFs rely on pre-existing open chromatin to bind their motifs. By contrast, at co-bound sites where BANP, GABPA, and ZFP143 do not affect open chromatin, there is minimal change in binding. To determine whether this dependency was unidirectional, we performed the reciprocal experiment: depletion of YY1 or MYC followed by ChIP-seq for BANP. As predicted, the absence of YY1 or MYC had no effect on BANP binding at shared CGI promoters (Extended data Fig.5d-f). Interestingly, we also observed interdependencies among the pioneer-like TFs themselves, where the removal of the dominant chromatin opener can also result in the loss of binding of the others.

**Figure 5.**
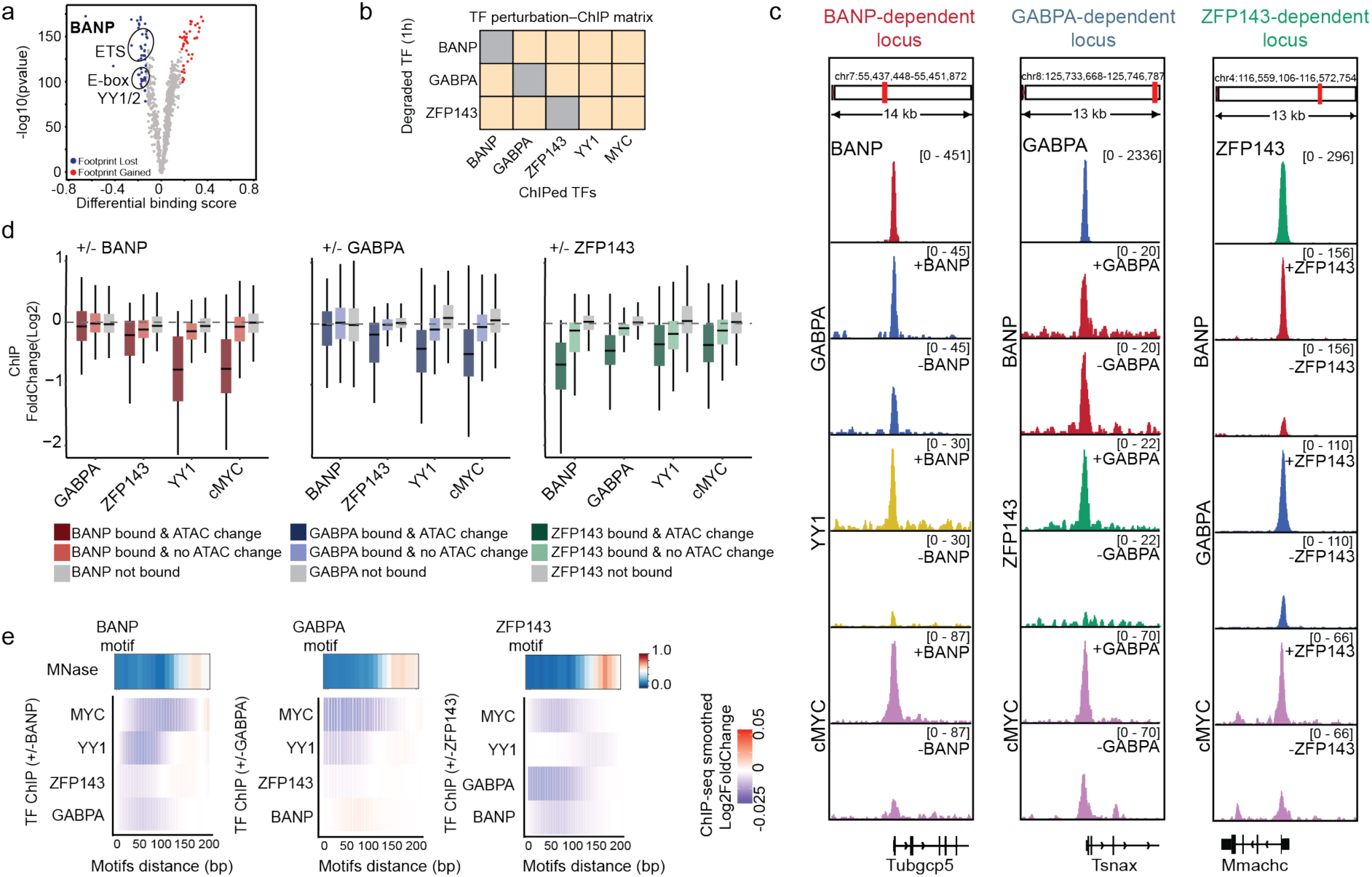
TF specific chromatin organization shapes co-binding hierarchies. a) Volcano plot displaying differential TF motif occupancy upon BANP degradation, as assessed by TOBIAS footprinting. Binding scores are plotted for individual motifs, highlighting motifs that show significant changes in occupancy following BANP loss. b) Schematic overview of the experimental approach to assess TF binding dependencies. Pioneer transcription factors were selectively degraded, and ChIP-seq was performed for other TFs to determine dependency relationships. c) A representative single locus example (Tubgcp5 promoter) illustrating BANP-dependent binding of co-bound TFs. d) The same single locus example as in (c), illustrating that BANP binding is independent of co-bound TFs. e) Dot plot depicting log₂ fold-changes in TF binding upon degradation of a pioneer factor. Only peaks that overlap between the degraded TF and the profiled TF are shown. Dot color indicates the specific TF. f) Heat maps showing the relationship between log2 fold-change in TF binding and the distance between co-bound motifs. Distances were binned in 50 bp intervals.

This exposes a hierarchical network of TF binding dependencies at co-bound promoters, where the ability of one factor to open chromatin enables binding of the additional factors. Given that open chromatin is surrounded by phased nucleosomes and that TFs interact with nucleosomes in different ways (REFs), these dependencies could be dictated by nucleosome position.

### Nucleosome position shapes binding hierarchies at CGI promoters

TF binding occurs in the context of chromatin, where nucleosomes restrict motif access^12^. Thus, the strong positioning of nucleosomes in proximity to sites bound by TFs such as BANP, GABPA, and ZFP143, as seen by MNase-seq, could shape the co-binding of additional TFs (Figure 5e, top track)^48^. To test this, we anchored our analysis on BANP, GABPA, or ZFP143 motifs and stratified the change in occupancy of co-bound TFs by the distance between motifs following the acute depletion of each anchor factor. This showed that loss of binding occurred most strongly 20-100 bp from the anchor TF, coinciding with the nucleosome-depleted region (Figure 5e). By contrast, there was a modest gain in binding very close (<20 bp), consistent with competitive occupancy, or further away (100-200 bp) within the nucleosome-occupied region, reflecting binding redistribution after anchor TF removal. These patterns indicate that nucleosome position modulates TF co-binding, with cooperative or competitive effects determined by the spatial relationship between motifs and nucleosomes. Together, we demonstrate that TFs cooperate at the level of binding and opening chromatin at a small number of sites, but does it also impact transcription?

### Cooperative activity occurs at suboptimal binding sites

To determine how TF co-binding translates into transcriptional regulation in the genome, we examined gene expression changes following individual and combinatorial TF depletion in our double-degron cell lines (Figure 6a,b, Extended data Fig.6a,c). First, to ensure that the data obtained using different degron tags are comparable, we confirmed that BANP depletion using either the dTAG or BromoTag system produced highly consistent results (Pearson correlation >0.77) (Extended data Fig.6b). We then asked whether simultaneous removal of two TFs produced transcriptional effects beyond those observed following single-factor loss. To quantify cooperative effects, we applied the interaction contrast design in DESeq2 to our data^49^, which distinguishes additive (sum of individual effects) from synergistic (greater-than-additive) responses. Consistent with our earlier findings, most co-bound genes were regulated by individual TFs (Extended data Fig.6d). In addition, a smaller subset of genes became significantly differentially expressed only after combined TF depletion (Figure 6c-e, Extended data Fig.6e,f). In most cases, the combined effect was additive, with true synergy being rare and confined to a handful of loci (Figure 6d,e, Extended data Fig.6f). Notably, the BANP-GABPA pair showed antagonistic interactions, where depletion of each factor alone had opposing transcriptional effects, with dual depletion yielding an intermediate expression level (Figure 6e).

**Figure 6.**
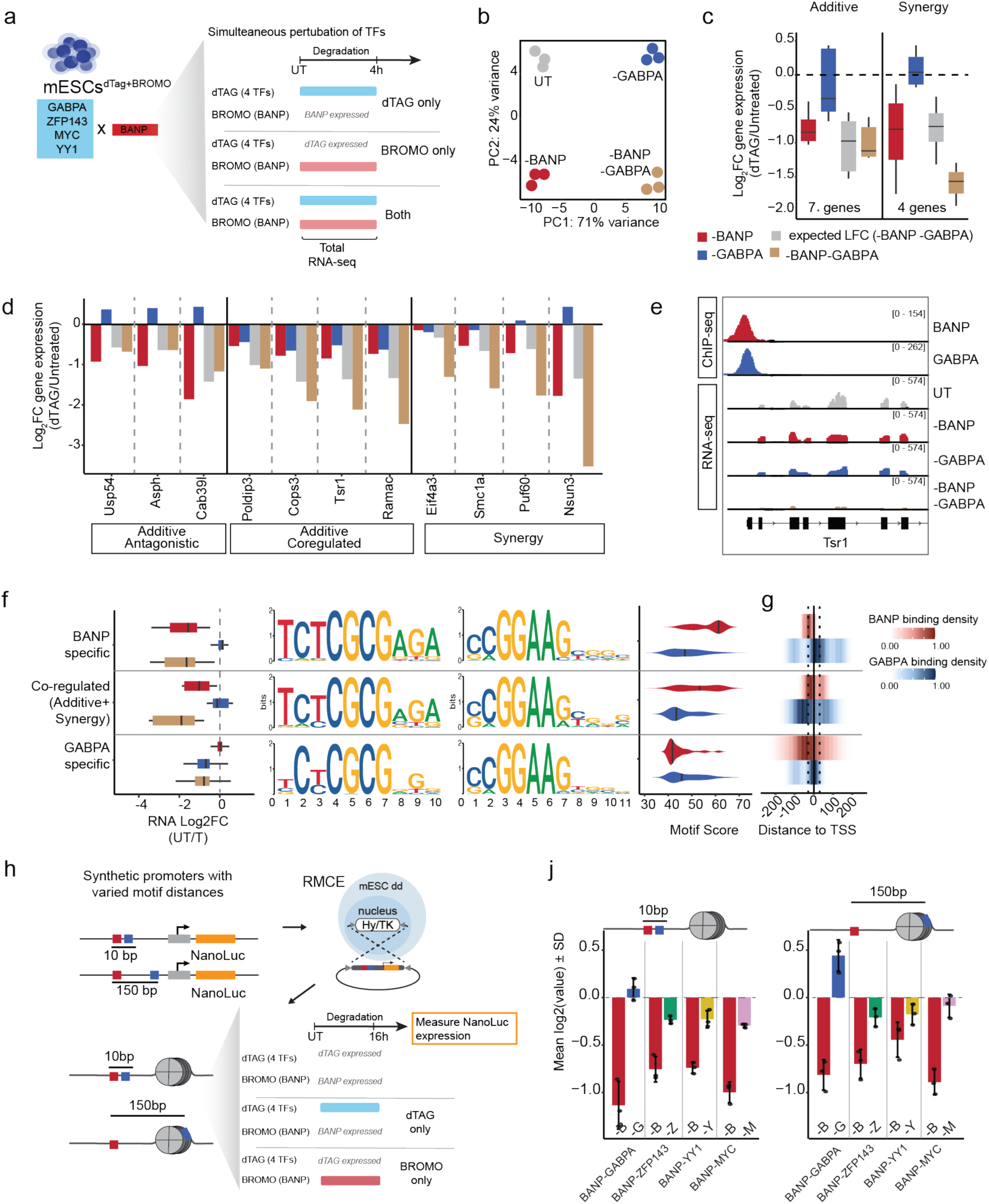
Co-binding of TFs can tune gene activity across cell types. a) Schematic of the experimental design used to assess combinatorial effects of dual TF depletion on gene expression. b) PCA of RNA-seq data showing transcriptional divergence after the degradation of individual or both TFs, relative to untreated (UT). c) Box plots showing log2 fold changes in gene expression upon individual or combined TF degradation for co-regulated genes identified by interaction contrast analysis. Observed changes are compared to expected additive effects (sum of single TF removals). Gene classes (additive or synergistic) were defined by interaction contrast analysis. d) Bar plots showing gene expression changes (log2 fold change) upon single or double TF degradation for all genes detected by interaction contrast analysis. Genes are grouped into three categories based on interaction contrast: additive-antagonistic, additive-co-regulated, and synergistic. e) Genome browser view of a representative locus illustrating a synergistically regulated gene. f) Motif variants of BANP and GABPA enriched at BANP-specific, coregulated and GABPA-specific promoters co-bound by the two factors. Violin plots showing the distribution of relative motif scores for each category. g) Density plot representing the distribution of BANP and GABPA binding. For each category, we generated a smoothed distribution of binding positions by computing a univariate KDE of distances from the TSS (−250 to +250 bp) with 200 evaluation points. Vertical reference lines were drawn at the TSS (0 bp; solid) and at ±30 bp (dotted). h) Schematic showing the design and targeted RMCE integration of synthetic promoters. j) Bar plot showing log2 fold-change in NanoLuc reporter expression of synthetic promoters upon the removal of BANP or a co-bound TF. TF motifs were positioned in the nucleosome-depleted (10bp) or nucleosome-occupied (150 bp) region, revealing distance-dependent effects on transcriptional output.

To explore what distinguishes promoters where a TF acts independently from those requiring cooperation, we examined motif variants, binding position relative to the TSS, and the distribution of co-bound TFs. We found that TF-specific promoters were enriched for the optimal motif of the dominant TF, whereas the accompanying factor typically occupied a suboptimal site (Figure 6f). In contrast, at cooperatively regulated sites, both factors engaged suboptimal motifs, suggesting that collaboration can compensate for weaker sequence recognition. Positional analysis further revealed that independent regulation coincided with binding of the dominant TF at its optimal distance from the TSS (as defined in Figure 3), while the partner TF was displaced from its preferred position. By comparison, co-regulated promoters were characterized by both factors deviating from the respective optimal binding position, an arrangement that may facilitate cooperative chromatin remodeling and transcriptional activation (Figure 6f). This shows that some co-bound TFs can have synergistic or antagonistic effects that fine-tune promoter output, potentially dictated by how TFs bind together depending on their position relative to a nucleosome.

### Synthetic reconstitution of TF dominance

Interpretation of our results in the genome is complicated by the presence of many motifs and co-binding events that we cannot account for (Figure 1b). To quantitatively assess the individual contribution of pairs of motifs to gene expression and how motif position relative to a nucleosome affects transcriptional output, we employed a synthetic system. First, we synthesised CGI-like promoters (500 bp long) with pairs of optimal motifs (BANP/GABPA, BANP/ZFP143, BANP/YY1, and BANP/MYC) placed 10 bp apart. These synthetic promoters were linked to a NanoLuc reporter and integrated separately into a defined genomic region in the corresponding double degron cell line using recombinase-mediated cassette exchange (RMCE)^50^. To quantify the impact each TF has on gene activity, they were individually degraded (16 h), and the NanoLuc activity was measured. The removal of BANP led to the strongest reduction in reporter activity across all constructs, comparable to the average loss in gene activity in the genome (Figure 2c and Figure 6g,h). The removal of the other TFs either did not affect (e.g., GABPA) or positively contributed to gene activity (e.g., ZFP143, YY1, and MYC), but only mildly in comparison to BANP. This indicates that BANP acts as the dominant TF in these promoters, despite being in the same sequence and chromatin context as GABPA and ZFP143, with additional factors not affecting or enhancing gene output. Interestingly, the cooperative action is strongly dampened compared to what was observed in transiently transfected plasmids (Figure 1d), suggesting that chromatin impacts cooperation.

Next, we tested whether the competition between TF binding and nucleosome occupancy could impact cooperative behavior. Since our synthetic promoters are dominated by BANP, which is known to strongly position nucleosomes (Figure 5e), we wondered if spacing the motifs at a nucleosomal distance (150 bp apart) would alter cooperative interactions^12,27^. Interestingly, when the additional motifs were placed in the nucleosome-occupied region, they antagonized (e.g., GABPA) or no longer contributed (e.g., MYC) to gene activity. This is in line with our interaction contrast analysis and implies that the organization of chromatin by dominant TFs results in a competition between additional TF binding and nucleosome occupancy, giving rise to chromatin-mediated activity modulation. These results show that strong co-activators can have context-dependent impacts on gene activity that are dictated by nucleosome positioning. Interestingly, the impact on gene expression for both ZFP143 and YY1 was not influenced by the distance between the motifs, however, placing YY1 at a nucleosomal distance, dampened the ability of BANP to activate expression. These results demonstrate that only one TF can be dominant at a specific locus, with equally capable TFs unable to compensate for removal. Furthermore, it suggests that the co-binding of two activators can synergistically or antagonistically modulate gene expression depending on their binding relative to a nucleosome and sensitivity to chromatin. This fits with our genomic observations of binding redistribution upon the degradation of a chromatin opening TF and that the depletion of an activator TF can also result in some genes being mildly upregulated.

Collectively, our results demonstrate that co-binding infrequently translates into the cooperative maintenance of promoter accessibility or transcriptional output and depends on promoter syntax. This suggests that TF co-occupancy often reflects structural or chromatin-level interactions rather than functional cooperation. Instead, our findings reveal that TF binding patterns at CGI promoters are shaped by one dominant TF that autonomously opens chromatin and drives transcription at distinct promoters, including CEGs. We further demonstrate a chromatin mediated mechanism whereby co-bound activators can tune gene expression synergistically as well as antagonistically. The inability of equally capable master TFs to compensate for the removal of the dominant TF at a specific locus and the fundamental requirement for CEGs to be expressed across tissues, would suggest that this mechanism is conserved.

### Conservation of regulatory dominance in human cells

Our experiments in mESCs revealed that chromatin accessibility and transcription at CGI promoters, including CEGs, are typically governed by single TFs. Given that these TFs drive the expression of essential genes, we reasoned that their function is likely conserved in humans, and that this should be reflected in mutational signatures. To explore this, we analysed naturally occurring non-coding variants in the human population from the gnomAD database and quantified their frequency within TF motifs located at promoters of essential versus non-essential genes^51^. The underlying assumption is that variants disrupting essential regulatory sites are depleted from the population due to their deleterious impact. Consistent with this, we found that TF motifs at essential gene promoters are significantly less frequently mutated than those at non-essential genes, suggesting strong purifying selection (Figure 7a).

**Figure 7.**
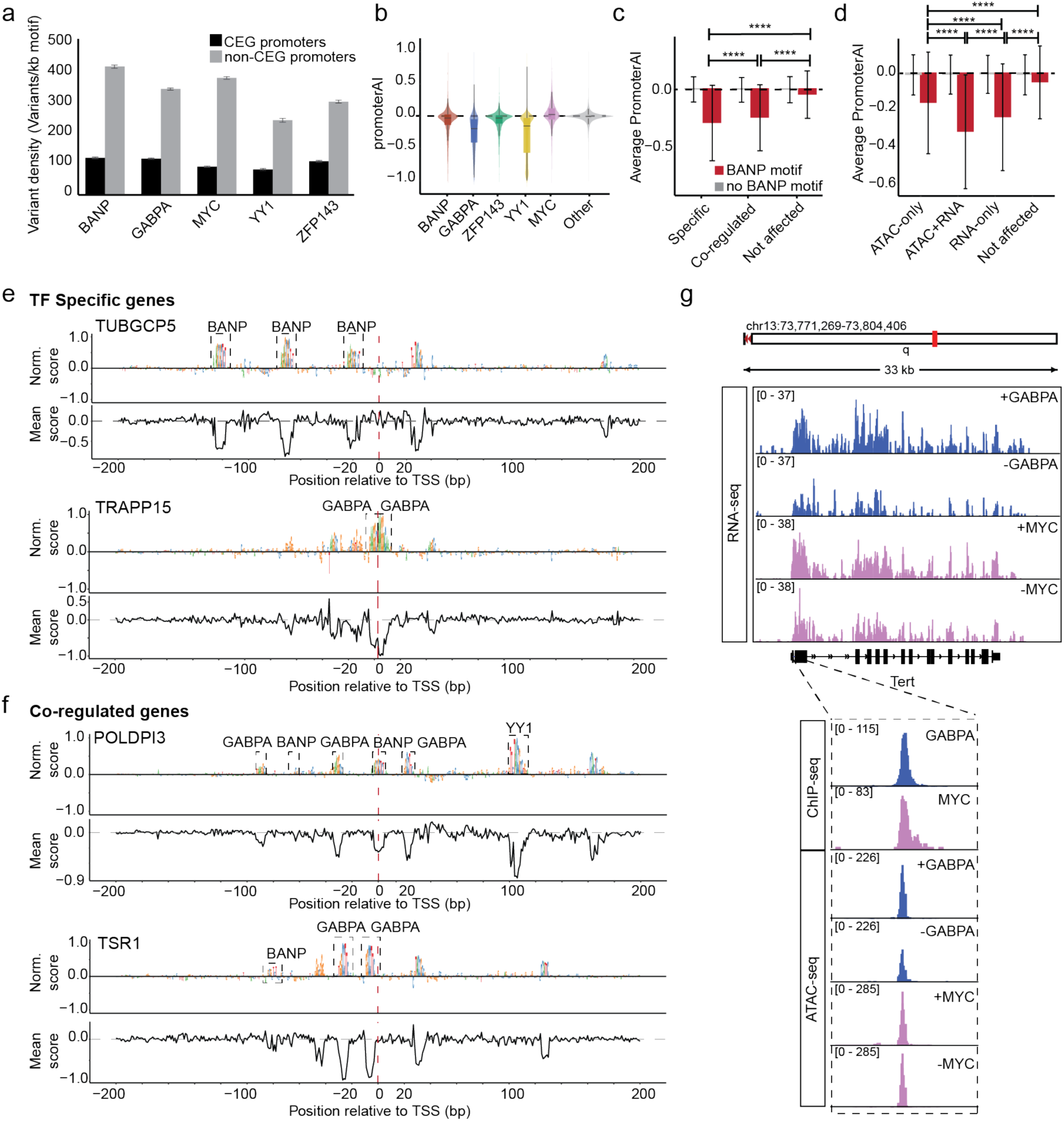
Conservation of regulatory dominance in human and variant effects. a) The frequency of motif variants found in BANP, GABPA, MYC, YY1, and ZFP143 sites in essential compared to non-essential genes. b) Violin plot showing the distribution of promoterAI scores (predicted variant effects) at the respective transcription factor (TF) motifs in CEG promoters in human. c) Average promoterAI score at the BANP motif for genes transcriptionally regulated exclusively by BANP, co-regulated, or not affected. Error bars indicate the standard error of the mean (SEM). d) Average promoterAI score at the BANP motif for genes regulated by BANP. Genes are categorized based on the type of regulatory effect: ATAC-only, ATAC+RNA, RNA-only, or not affected. Error bars represent SEM. e) Attribution maps for representative single-promoter-regulated genes (TUBGCP5, upregulated; TRAPPC15, downregulated). Each panel shows a 400 bp window centered on the transcription start site (TSS). The upper track displays the attribution map with normalized promoterAI scores, and the lower track shows a line plot representing the mean promoterAI score across the same region. f) Same as (e) but for promoters co-regulated by BANP and GABPA (POLDPI3 and TSR1). g) The impact on transcription and accessibility at the TERT gene upon degradation of each TF.

Given this, we wondered if signatures of regulatory dominance could be observed in human cells. To explore this, we leveraged precomputed PromoterAI scores, which are generated by a deep learning model that predicts the effect of all possible single-nucleotide variants in promoters of human protein-coding genes on gene expression^31^. This approach provides an orthogonal perspective to our degron-based perturbations, quantifying the predicted impact of motif disruption rather than TF depletion. First, we examined whether the motifs of the five TFs, BANP, GABPA, ZFP143, YY1, and MYC, are predicted to drive essential gene expression in human promoters. Indeed, promoterAI predicted that mutations within these motifs would lead to reduced CEG expression, especially when located proximal to the TSS, compared to mutations in random 10 bp control sequences within the same promoters (Figure 7b, Extended data Fig.7a). The only exception was MYC, whose motif mutations had minimal predicted effect, consistent with our experimental observations that MYC functions primarily as a transcriptional amplifier and not a dominant driver^52^. Interestingly, promoterAI also predicted stronger downregulation of genes upon the disruption of GABPA, ZFP143, and YY1 motifs at promoters that were unaffected by TF depletion. Given that promoterAI was trained on many tissue types, this could reflect that BANP function is highly conserved across species and tissues, while GABPA, ZFP143, and YY1 activity is more varied. It could also be a result of the different types of perturbations being made: promoterAI simulates motif disruption, potentially affecting all TFs capable of recognizing that sequence, whereas degron experiments isolate the effect of removing a single TF. Given that GABPA belongs to the large ETS family of TFs, and ZFP143 and YY1 share motifs with related zinc-finger proteins, motif disruption could reduce binding of multiple factors, resulting in a predicted impact not observed upon individual TF depletion. Conversely, BANP motifs, which are less promiscuous, may be recognized by fewer alternative TFs, yielding higher concordance between experimental and predicted effects.

Next, we asked whether the promoterAI-predicted variant effects recapitulate the different TF regulatory modes identified in our degron experiments. For each TF, we compared the average predicted variant effect across distinct gene classes derived from our data: TF-specific, co-regulated, and unaffected genes. Variants within BANP motifs showed the strongest predicted effects at TF-specific targets, weaker effects at co-regulated genes, and negligible effects at unaffected promoters (Figure 7c). This was less clear for the other factors, with similar effects for all groups (Extended data Fig.7b). Moreover, stratifying the genes by ATAC-only, RNA-only, or both revealed that variants were most detrimental at loci where the TF acts on both chromatin and transcription, consistent with our experimental hierarchy (Figure 7d). Single locus examples of TF-specific targets display strong predicted sensitivity to variants only within BANP or GABPA motifs clustered near the TSS, despite extensive co-binding of additional TFs (Figure 7e, Extended data Fig.7c). By contrast, at BANP and GABPA co-regulated targets, combinations of motifs are predicted to impact gene expression (Figure 7d,e). Interestingly, the impact of motif variants also scales with our observations, with stronger contributions of individual motifs at the specific target compared to the co-regulated ones (Figure 7d,e). This illustrates how a single nucleotide change can quantitatively modulate the expression of an essential gene through altering TF occupancy.

These analyses demonstrate that the hierarchical regulatory logic uncovered in mouse is conserved across species and reflected in human promoter architecture. They also highlight the complementarity between experimental perturbations and computational variant prediction.

### TF hierarchies explain predicted effects of promoter variants

Non-coding genetic variants are often associated with disease, but a mechanistic understanding of how this impacts gene expression is often unclear. Non-coding somatic mutations in the TERT gene are some of the most common in human cancer cells^53^. Two highly recurrent mutation hotspots in the TERT promoter have been shown to create novel consensus binding sites for ETS family TFs, leading to upregulated telomerase expression and decreased cell death^31,54^. In addition, MYC is often overexpressed in cancer cells, which is linked to increased TERT gene expression^55^. However, the underlying mechanisms of how these features collectively modulate TERT expression remain unclear. We now provide experimental evidence that confirms that both GABPA and MYC contribute to the expression of the TERT gene (Figure 7g). Our experiments also show that GABPA is the dominant factor at the TERT promoter, being able to drive the opening of chromatin that likely facilitates the co-binding of MYC (Figure 7g). The co-binding of MYC contributes to expression in a synergistic way together with GABPA (Figure 7g). These mechanistic insights allow us to explain how combining the gain in ETS motifs, which likely results in GABPA being able to increase chromatin accessibility, and the overexpression of MYC would result in increased expression of the TERT gene. These results demonstrated that defining the regulatory hierarchies and combinatorial contributions to gene regulation enables to better understand how gene expression is driven and altered in disease.

## Discussion

The regulatory networks and hierarchies that ensure the expression of CEGs to sustain cell viability have been challenging to study and remain unresolved. Universal open chromatin and dense TF binding patterns at the CGI promoters that control CEG expression have led to the common interpretation that robust transcription is achieved by cooperative and redundant TF action^53,56,57^. Furthermore, it is unclear how different cell types tune the expression of CEGs to adapt to cellular requirements. Here, we demonstrate that individual TFs drive the ubiquitous expression of distinct sets of CGI promoter-controlled genes, including CEG, by dominantly opening chromatin and activating transcription. These master TFs control the expression of key components of core cellular machinery, with their disruption resulting in cell death. The observation that CEG expression level alone can distinguish cell types indicated the need for tight control and spurred us to nominate key TFs that regulate their expression. The systematic dissection of the top five-ranked critical TFs using individual and combinatorial inducible degradation and recovery captured direct primary targets, enabling us to define regulatory hierarchies that govern CGI activity. Particularly, our data uncover a previously unrecognized regulatory architecture where CGI promoters are dominantly controlled by single master TFs that continually establish active chromatin (accessibility and histone modifications) and transcription at distinct sites. This regulatory dominance was reconstructed in synthetic promoters and was mirrored in motif variant data in humans, showing evolutionary conservation, potentially as far back as zebrafish, where BANP controls similar genes^58^. Furthermore, the conserved action of these TFs across cell types and during human development suggests that this is a generalizable mechanism^30^. Dominant activity is underpinned by the ability to autonomously open chromatin, which is facilitated by the presence of a strong motif and binding, and translates into productive transcription when located proximal to a TSS. These features predict that TFs able to open chromatin and found enriched close to the TSS at promoters (e.g., NRF1, NFY, and SP/KLF factors), would also be CGI master regulators^12,30,39^. Indeed, this has been seen for the pioneer factors Oct4::Sox2 and the CGI binder NRF1^59–61^. Of note, while the YY1 motif has been associated with universal open chromatin^30^, we do not see a strong effect on chromatin accessibility upon its removal, hinting at potential redundancy with the related TF YY2^62^. Collectively, this reveals that the robust encoding of cell viability is achieved by ensuring the proper functionality of a single TF, rather than complex networks of TFs. While efficient, this makes the systems highly sensitive to mutations that disrupt these TF, with failure of the dominant factor resulting in cell death, acting as a checkpoint for core cellular functionality.

A pivotal concept In development biology over the past decades has been the idea of pioneer TFs that can bind to nucleosome-occupied DNA and initiate remodeling to make regulatory regions accessible for other factors. They act as““pionee”” organizers of the chromatin landscape, setting the stage for subsequent gene activation during development and cell fate decisions^13,63^. By contrast, CGI promoters that drive ubiquitous gene expression typically exhibit constitutively open chromatin and are occupied by dense TF binding, indicative of highly interconnected regulatory networks^64^. In stark contrast, our data demonstrates that individual TFs are necessary and sufficient to drive the continual opening of chromatin at CGI promoters. Interestingly, our combinatorial degron and recovery experiments show that pioneer activity is highly locus-specific and mostly non-redundant, with co-occupancy of multiple chromatin-opening TFs not able to compensate for the removal of the dominant factor. This suggests that pioneer activity is context-dependent, which is exemplified by factors such as GABPA and NRF1 not being able to open chromatin in a synthetic context, yet being able to pioneer specific loci in the genome^1265^. This might indicate that many TFs have the potential to pioneer loci at the right time and place^15^, potentially through the recruitment of specific cofactors^66^. Furthermore, we unequivocally show that the extensive co-binding of additional TFs is enabled by the chromatin-opener TF and is shaped by nucleosome position. This unidirectional dependence at co-bound promoters has also been observed in yeast^67^. While co-binding infrequently resulted in co-regulation of the linked gene, both antagonistic and synergistic impacts were observed at cooperative sites, likely dependent on the interplay with chromatin, as seen with synthetic promoters. Such a hierarchical organization simplifies the complex regulatory networks currently thought to control essential genes, with dependence on a single chromatin-insensitive TF to ensure expression. The co-binding of additional TFs, including cell-type-specific TFs, can modulate the output through chromatin-mediated cooperativity, potentially enabling cells to adapt to cellular demands^14,15,66^. Thus, we define and put forward a pioneer model for CGI promoters, where individual dominant TFs are necessary and sufficient to open chromatin and drive transcription.

A hallmark of CGI promoters is the presence of active histone modifications, such as H3K4me3 and H3K27ac^68,69^. Yet, their dynamics and link to chromatin organization and transcription remain unclear. Profiling the immediate changes in active chromatin marks found that only genes that lost open chromatin and transcriptional activity upon BANP removal showed a concurrent reorganization of H3K4me3 and loss of H3K27ac. This would suggest that the presence of these marks is not sufficient for transcription, and that these marks are not directly involved in opening chromatin or activating genes. This is in line with work that showed little effect on gene expression upon the removal of acetylation at lysine 27 on histone H3 by alanine substitution, ablating the presence of the H3K27ac mark^70^, or loss of the major H3K4me3 modifier SET1^71^. While forced addition of H3K27ac has been shown to stimulate gene activation^72^, recent work with reconstituted systems suggested that this could occur by controlling chromatin architecture^73^. Interestingly, the strong link with BANP could indicate that it is able to recruit the enzymes that place these marks directly, while at genes regulated by the other TFs, they are recruited by another TF.

These findings demonstrate that the ubiquitous expression of genes is not ensured by complex redundant regulatory networks, but rather by a simple hierarchy that is dominated by one TF, which represents a paradigm-shift in our understanding of CGI promoter activity. While rare, TFs can cooperate at two levels: 1) where chromatin accessibility is established by one TF, but robust transcription requires additional TFs that enhance RNA output, and 2) where two or more TFs act jointly to open chromatin and drive transcription, representing fully collaborative activation. These cooperative modes likely confer additional plasticity, allowing modulation of gene expression in specific cellular or developmental contexts to adapt to cellular requirements. Beyond essential genes, these findings offer a paradigm for how regulatory element organization can encode regulatory hierarchy across the genome. Indeed, soft motif syntax between suboptimal motifs has been shown to support cooperation also at enhancers when within a nucleosomal distance^74^. Furthermore, it provides a foundation for interpreting the functional impact of noncoding variation and guiding efforts to model gene regulation in health and disease.

Developmental and disease research has been largely focused on defining the cell-type-specific factors that drive these processes, with little attention given to changes in CEG regulation. With the advancement in deep learning-based models, it has become evident that motifs recognized by ubiquitously expressed TFs also play key roles in development and disease^30,31^. Our work shows the dominant role some of these ubiquitously expressed TFs have in establishing and maintaining gene activity of core essential genes, underscoring their important role in these processes. This also supports the recently shown critical role for GABPA in zygotic genome activation (ZGA)^36^, and would suggest that factors like BANP and ZFP143 could also play an important role in ZGA. Furthermore, it is becoming clear that the misregulation of essential cellular processes can contribute to healthy aging and disease. For example, a key feature of aging is the downregulation of essential genes^3^. Alternatively, the overexpression of the essential TF cMYC is associated with many cancer types^4^. Therefore, defining the mechanistic basis of CEG regulation provides a framework for therapeutic strategies aimed at restoring or modulating essential gene activity.

Finally, the rapid expansion of deep learning models capable of extracting regulatory features from genomic and epigenomic data is transforming our ability to predict gene expression^31,75,76^. Our results align with promoterAI predictions of the impact of motif variants on promoter activity, yet discrepancies between predicted and observed effects underscore complementarity of the approaches and the importance of experimental data. This is especially critical for interpreting disease-associated noncoding variants, where identifying the correct TF effector is essential for precision therapeutics. By defining the regulatory dependencies that govern CGI promoters, our work provides a foundation for leveraging both experimental and computational approaches to predict, manipulate, and restore essential gene expression in development, aging, and disease.

## Data availability

Next-generation sequencing data are deposited at the Gene Expression Omnibus database.

## Acknowledgments

R.S.G. is funded by a DFG grant (GR 6341/2-1). We thank Paul Ginno, Luke Isbel, Anaïs Bardet, and members of the R.S.G. laboratory for critical feedback on the manuscript.

## Author contributions

R.S.G. and M.C. conceived and planned the experiments. M.C. performed the majority of experiments and did the computational analysis. S.N. performed RMCE and NanoLuc assays, helped with cloning, cell line generation, validation, and maintenance. I.S.A. helped with ChIP-seq and ATAC-seq experiments. S.D. helped with cell line generation, validation, and maintenance. F.B.T. performed single-cell analysis with guidance from M.C. and R.S.G. R.S.G. supervised the project. R.S.G. and M.C. interpreted the results and wrote the manuscript.

## Methods

### – Cell culture

#### ES cells

TC-1 mouse embryonic stem cells (129S6/SvEvTac background; originally provided by A. Dean, NIH) carrying an RMCE site within the γ-globin locus served as the wild-type line. Cells were cultured in Dulbecco’s Modified Eagle Medium (DMEM; Invitrogen) supplemented with 15% fetal calf serum (Invitrogen), L-glutamine (Gibco), and non-essential amino acids (Gibco). The medium was further enriched with β-mercaptoethanol (Sigma) and leukaemia inhibitory factor (LIF) produced in-house. For all assays, cells were expanded over several passages on 0.2% gelatin-coated plates (Sigma).

### – Generation of cell lines

#### Single degron

For inducible GABPA, ZFP143, YY1 and cMYC depletion in wild-type mouse ES cells, we applied the FKBP12(F36V)-based dTAG system as previously described by Grand et al. Briefly, each TF was endogenously tagged at the C-terminus portion with FKBP12(F36V)-V5 using CRISPR/Cas9-mediated knock-in with gRNA and repare-template plasmids, followed by puromycin selection, clonal isolation, and genotyping. Correctly targeted clones were validated by PCR, Sanger sequencing and western blotting before and after dTAG13 treatment, using vinculin as a loading control.

#### Double degron

For dual protein depletion, single degron ES cell lines were re-engineered using CRISPR/Cas9-mediated knock-in to insert an HA-Bromo tag at the C-terminus of the Banp gene, following the strategy established for the single degron system. The functionality of the additional degron was verified by western blotting after treatment with the corresponding small molecules (dTAG13 for FKBP12(F36V) and AGB1 for the Bromo tag).

### – Recombinase-mediated cassette exchange (RMCE) for targeted integration in mouse ES cells

RMCE was performed as previously described (21964573). In brief, double-degron modified TC-1 ES cells (background 129S6/SvEvTac) carrying an RMCE selection cassette were selected under hygromycin (250 μg/ml, Roche) for 10 days. Next, 0.5 million cells were transfected with 1 μg of reporter plasmid and 750 ng of pIC-Cre plasmid using Lipofectamine 3000 Reagent (Thermo Fisher Scientific). Negative selection with 3 μM Ganciclovir (Roche) was started 2 days after transfection and continued for 10 days. Selected cells were plated for clone picking, and individual clones were PCR screened to ensure the insertion of the reporter constructs all in the same orientation.

### – Cell death assay

TF-dTAG cell lines were seeded in triplicate into gelatin-coated 6-well plates at a density of 5 × 10⁵ cells per well. After 24 h, triplicate wells of each line were treated with dTAG13 (500 nM; Sigma). Viable cells from both untreated and treated conditions were counted every 24 h using a TC20 automatic cell counter (Bio-Rad).

### – Luciferase assay

#### Transient transfection

Single, dual, and pairwise combinations of TF motifs or scrambled control sequences, separated by a 5-bp spacer, were inserted into the centre of a GCI-like sequence and cloned upstream of a NanoLuciferase reporter gene. The nanoluciferase reporter plasmid (250 ng) and Renilla firefly luciferase reporter plasmid (25 ng) were co-transfected into 50,000 TC-1 ES cells in suspension using Lipofectamine 3000 Reagent (Thermo Fisher Scientific) in a 96-well plate. After 24 hours, the reporter gene activity was measured using the Nano-Glo Dual-Luciferase Report Assay System (Promega) according to the manufacturer’s instructions, and the fold increase in activity of the constructs relative to the scrambled control was calculated for three independent biological replicates.

#### Stable cell lines

Pairwise combinations of the BANP motif with all other tested TF motifs (GABPA, ZFP143, YY1, and MYC), 10 bp apart or 150 bp apart, were inserted into the GCI-like promoter sequence and cloned upstream of a NanoLuciferase reporter gene. These constructs were stably inserted into the genome of the corresponding double-degron cell line by RMCE as described above (see ‘Recombinase-mediated cassette exchange (RMCE) for targeted integration in mouse ES cells’). After selection, single clones were picked and screened for construct insertion and orientation, and three clones for each construct were expanded for the luciferase assay. The day before the experiment, each cell line and clone (1 million cells) was transfected with a Renilla firefly luciferase reporter plasmid (500 ng) in suspension using Lipofectamine 3000 Reagent (Thermo Fisher Scientific). Then, 60,000 cells were plated in triplicate for each condition into a 96-well plate. The following day, the cells were treated with with the dTAG or the BromoTag for 16h and NanoLuc activity was measured using the Nano-Glo Dual-Luciferase Report Assay System (Promega) according to the manufacturer’s instructions. The fold change in activity relative to the untreated control samples was calculated for three independent biological replicates.

### – ChIP-seq

TF-dTAG cells (1.2 × 10⁶) were plated on 15-cm dishes one day before the experiment. ChIP was performed as previously described^77^ with the following modifications: chromatin was sonicated for 20 cycles of 20 s on/40 s off using a Diagenode Bioruptor Pico; 75–200 µg of chromatin was used per immunoprecipitation. Immunoprecipitated DNA was used for library preparation with the NEBNext Ultra II DNA Library Prep Kit (Illumina), including 12 PCR cycles for amplification. Libraries were sequenced on an Illumina NextSeq2000 P2 100bp. To test the effect of TF degradation on binding or histone mark levels, two plates per replicate were prepared. One hour before ChIP, the medium in all plates was exchanged, and dTAG13 (500 nM; Sigma) was added to half of them to induce degradation of the targeted TF.

### – RNA-seq

TF-dTAG and TF-double degron cells (2.5 × 10⁵) were seeded in triplicate into 6-well plates one day prior to the experiment. To induce TF degradation, the medium was replaced with fresh medium containing the appropriate compound (500 nM; Sigma) for the indicated duration (1 h or 4 h). For simultaneous depletion of both proteins, both compounds were added together. Total RNA was extracted using the Direct-zol RNA Purification Kit (Zymo) according to the manufacturer’s instructions. Sequencing libraries were generated from 100 ng of total RNA (three biological replicates) using the TruSeq Stranded Total RNA Library Prep Kit (Illumina). Libraries were sequenced on an Illumina HiSeq platform (50 cycles).

### – ATAC-seq

TF-dTAG and TF-double degron cells (2 × 10⁵) per replicate were plated on a 6-well plate the day before the assay. Each replicate was plated twice. 1h before starting the protocol, the media was changed in all the wells and in half of them the specific TF degradation was induced upon addition of the specific compund to the media.

The day after, 7.5 × 10⁴ cells were harvested for the ATAC-seq experiment, and the leftover was used to run total protein extract and verify with westernblotting the correct total TF depletion in the treated samples.

ATAC-seq was performed using pre-indexed assembled Tn5 transposomes (Active motif, https://www.activemotif.com/documents/2267.pdf) according to the manufacturer’s instructions with minor modifications. Briefly, 7.5 × 10⁴ cells were harvested, washed twice with ice-cold PBS, and lysed in ATAC lysis buffer. Nuclei were pelleted and resuspended in tagmentation mix (25 µl 2× Tagmentation buffer, 2 µl 10× PBS, 0.5 µl 1% digitonin, 0.5 µl 10% Tween-20, and 18 µl nuclease-free water). Each sample was combined with pre-indexed Tn5 and incubated for 30 min at 37 °C with shaking at 800 rpm. Reactions were stopped with 1.8 µl 0.5 M EDTA and 0.5 µl 10% SDS, followed by incubation at 55 °C for 3 min. DNA was purified with AMPure XP beads at a 1.2× ratio and eluted in 22 µl TE buffer.

Libraries were generated from 20 µl of eluted DNA using NEBNext Ultra II Q5 Master Mix (Illumina kit) and 1.25 µM P5/P7 primers, with the following PCR program: 72 °C for 5 min; 98 °C for 30 s; 8 cycles of 98 °C for 10 s, 63 °C for 30 s, and 72 °C for 1 min; final hold at 4 °C. PCR products were purified using a 1.2× AMPure cleanup and eluted in 17 µl TE buffer.

Indexed samples were quantified and normalized to 6 nM, pooled, and further purified by a double-sided AMPure bead selection (0.6× followed by 0.9×) according to Illumina guidelines (REF). Final pools were eluted in nuclease-free water, quantified by Qubit and fragment distribution was verified by bionaliser. Sequencing was carried out on the Illumina HiSeq platform (50 cycles).

### – TF recovery

For experiments involving recovery condition, cells were first treated with the compound inducing TF degradation for 4 h. Subsequently, cells were trypsinized, collected, and washed three times with 1× PBS by inverting the tube 5–10 times followed by centrifugation (3 min, 1000 rpm). After the final wash, cells were resuspended in the original volume of fresh medium lacking the compound and re-plated. Recovery conditions were assessed 24 h after re-plating.

## Methods: Computational Analyses

### – Annotations

Promoters were defined as -2000 +200 nt around the TSS of each transcript in the UCSC Known Genes database, which was accessed via the Bioconductor packages TxDb.Mmusculus.UCSC.mm39.knownGene (v.3.10.0) for mouse.

CpG islands were retrieved from UCSC Genome Browser (mm39 assembly) (‘cpgIslandExt’ table) using rtracklayer, and converted to GRanges objects. Promoters overlapping CpG islands were classified as CpG island promoters, while the remainder were assigned as non-CpG island promoters.

Enhancer-gene associations for mouse embryonic stem cells were obtained from publicly available enhancer atlas resources (EnhancerAtlas 2.0^78^).

Common Essential Genes were retrieved from DepMap, as results of CRISPR-KO screening.

Genome-wide transcription factor binding sites in mouse ESCs were obtained from the ReMap database^26^, restricting to ChIP-seq experiments performed in mESCs. Only factors with direct DNA-binding activity were retaine. The comprehensive list of mouse transcription factors was retrieved from the Mouse Transcription Factor Atlas^79^. As BANP was not present in the ReMap dataset, it was manually incorporated.

### – TFs enrichment analysis at CREs

In order to calculate the average number of TF binding events at different CREs, we first aimed to filter promoters for active genes only.

Promoters and enhancers were classified as “active” if the associated gene was expressed in mESC according to our RNA-seq in wild-type condition (TMP≥1). For downstream analyses, enhancers were filtered to include only regions between 1500-3000 bp in length, in order to make the length of the region comparable to the promoters. Genome-wide transcription factor binding sites in mouse ESCs were retrieved from the ReMap database. Binding site coordinates were converted into GRanges objects. Overlaps between TF binding sites and genomic regions of interest (CpG island promoters of expressed genes, non-CpG island promoters of expressed genes, and filtered enhancers) were quantified using the countOverlaps function.The mean number of TF binding events per region type was calculated and visualized by violin plots (ggplot2), comparing CpG island promoters, non-CpG island promoters, and enhancers. Statistical differences in TF binding between groups were tested by pairwise t-tests with Benjamini–Hochberg correction for multiple testing.

To quantify the diversity of transcription factor (TF) occupancy across regulatory elements, Shannon entropy was calculated for expressed CpG island promoters, non-CpG island promoters, and enhancers. For each region, a binary matrix was generated, marking presence/absence of TF binding per TF. Shannon entropy was computed per region as *pilog2pi*, where pi represents the relative frequency of TF binding events. Distributions of entropy across region types were visualized using density plots. All analyses were performed in R using the GenomicRanges, dplyr, and ggplot2 packages. This analysis provides a measure of TF binding diversity, where higher entropy values indicate a broader and more heterogeneous set of TFs bound to a region, while lower values reflect more specific TF occupancy.

### – Single-cell RNA-seq Analysis

Single-cell RNA-seq data were obtained from the FACS-based Tabula Muris brain dataset. Data preprocessing and analysis were performed in Seurat v5.0.2 (R v4.4.3). After standard quality control, UMI counts were normalized using LogNormalize(). Common essential genes (CEGs, defined by DepMap) were set as highly variable features (FindVariableFeatures()), and only these genes were used for downstream analysis. Data were scaled (ScaleData()), and PCA was performed (RunPCA()), retaining the top 15 of 50 components. Cells were clustered using a KNN graph-based method (FindNeighbors() and FindClusters(); resolution = 1). Differential expression was computed in a one-vs-all manner with FindAllMarkers(), using cell ontology classes as labels. Genes were considered differentially expressed if adjusted p < 0.5 and expressed in ≥50% of cells in both the focal and comparison groups, to minimize artifacts due to sparse detection. Cell-state visualization was performed using UMAP (RunUMAP(), dims = 15) computed on the CEGs. Plots were generated with ggplot2 (v3.5.1), with cells colored by ontology class..

### – ChIP-seq

#### Mapping and peak calling

Raw ChIP-seq reads were processed using SnakePipes (v2.8.0, GitHub: https://github.com/maxplanck-ie/snakepipes). Reads were mapped to the mouse genome (GRCm39) with adapter trimming, PCR duplicate removal, and a minimum mapping quality threshold of 10. ChIP-seq signal processing, including peak calling and quality control, was performed using the SnakePipes ChIP-seq workflow with MACS2, in single-end mode with a fragment length of 200 bp. Default parameters were used unless otherwise specified.

#### Consenus peak processing and annotation

ChIP-seq peaks from multiple replicates were processed using R and the GenomicRanges framework. BED files of individual replicates summit peaks were imported and converted into GRanges objects. Non-canonical chromosomes were removed, and peaks overlapping blacklisted genomic regions were excluded.

GRanges objects were resized to 200 bp centered on the peak summits. For each TF, a set of consensus peak summits were computed as the midpoint of overlapping peaks between replicates. Consensus peaks were annotated to genes using ChIPseeker with the annoatatePeak function and the TxDb.Mmusculus.UCSC.mm39.knownGene transcript database, defining promoters spanning -2kb to 200bp relative to the TSS. Binding intensity was quantified by counting reads overlapping each consensus peak for each replicate, using featureCounts. Counts were then normalized to counts per million (CPM) for each replicate, and the mean CPM across replicates was used as the representative binding intensity for each peak.

For each TF, the top 1000 consenus peaks with the highest read counts were selected. Consensus peak summits were resized 100 bp and the relative DNA sequences were extracted using Bsgenome.Mmusculus.UCSC.mm39 and exported in FASTA format for motif enrichment analysis with MEME suite. For each of the five transcription factors, the V5 ChIP was enriched for the expected motif. Since the BANP motif is not included in the JASPAR database, the top motif identified in the BANP ChIP was ZBTB33/Kaiso.

#### Merge set of peaks between TFs

To generate a unified set of binding sites across the five TFs, consensus peaks were resized to 200 bp centered on the summit, concatenated, sorted and merged using BEDTools to obtain a non-redundant set of genomic intervals. Because merged regions varied in size, new summits were recalculated as the midpoint of each interval, and windows of 200 bp centered on the summit were generated for downstream analyses. This resulted in a set of 29,288 peaks, classified into CpG promoters, non-CpG promoters, and distal elements. Each peak was further annotated for TF occupancy by assessing overlap with individual TF consensus peak sets.

### – RNA-seq

RNA-seq data were processed using the snakePipes mRNA-seq workflow. Raw sequencing reads in FASTQ format were aligned to the mouse reference genome (GRCm39) with default settings, including adapter and quality trimming. Gene-level quantification was performed with featureCounts, as implemented in the pipeline. The workflow generated alignment files, quality-control reports, and count matrices, which were subsequently used for downstream differential expression analysis.

To capture early transcriptional responses, we separately quantified intronic and exonic reads using eisaR (v1.18.0; Gaidatzis et al., 2015). Exonic and gene-body annotations were derived from Ensembl GTF files (release 106) with GenomicFeatures (v1.36.4). Counts were obtained with featureCounts, with intronic counts calculated as the difference between gene-body and exon counts. Genes with mean counts ≤ 5 were excluded. Differential expression analysis was then performed independently on intronic and exonic counts with DESeq2, enabling detection of rapid transcriptional changes at the 1 h and 4 h time points.

Because not all genes contain introns or yield detectable intronic signal, primary affected genes were defined as follows: if intronic changes were detectable, they were prioritized; otherwise, exon changes were considered. TF primary target genes were identified as genes significantly affected (padj < 0.01) at either the intronic or exon level at 4 h of TF degradation, provided they were also associated with binding of the corresponding transcription factor.

#### Interaction contrast

Differential expression analysis was performed on intronic RNA-seq counts to quantify the effects of single and combined transcription factor (TF) depletion. Count matrices were generated with featureCounts and processed with DESeq2 (v1.42.0). Experimental conditions were specified to reflect the independent or simultaneous removal of BANP and TF2, and a factorial design was applied with the formula

Design = ∼banp_removal x TF2_removal

This model estimates the main effects of BANP and TF2 depletion as well as their statistical interaction. Differential expression contrasts were extracted for: (i) BANP removal alone, (ii) TF2 removal alone, (iii) the interaction term (representing non-additive effects of double removal), and (iv) the combined effect of both TFs versus untreated control.

Genes were classified into regulatory categories based on adjusted p-values (padj < 0.05). Genes significantly affected by BANP removal only were labeled BANP-specific, while those affected only by GABPA removal were labeled TF2-specific. Genes significantly altered by both single perturbations but not by the interaction term were considered co-regulated. Genes with a significant interaction effect were classified as synthetic, indicating non-additive regulation under double TF depletion.

This classification enabled systematic dissection of TF-specific, co-regulated, and synthetic transcriptional responses.

### – ATACseq

ATAC-seq data were processed using the snakePipes workflow. Raw sequencing reads in FASTQ format were first preprocessed with the DNA-mapping module, which performs adapter trimming, PCR duplicate removal, and alignment to the mouse reference genome (GRCm39) using Bowtie2 with a minimum mapping quality threshold of 10. The resulting alignment files were subsequently processed with the ATAC-seq module of snakePipes, which generates genome-wide accessibility profiles, performs peak calling, and produces standardized quality-control reports. Peak calling was carried out with MACS2, and both fragment length distributions and transcription start site (TSS) enrichment scores were computed to assess data quality.

Default pipeline parameters were applied. The workflow produced alignment files, normalized coverage tracks, and set of accessible peaks for downstream analyses.

#### Differential Accessibility Analysis

For differential chromatin accessibility analysis following TF degradation, ATAC-seq reads were quantified over the unified set of merged peaks generated by combining the consensus peak sets of all TFs (as defined above). Read counts per peak were summarized into a count matrix and used as input for DESeq2 (v1.42.0). Differential accessibility was evaluated by comparing untreated (UT) samples with those collected 1 h and 4 h after TF depletion. Peaks with an adjusted P value (padj) < 0.05 were considered significantly regulated. Direct TF-dependent ATAC-seq targets were defined as significantly regulated peaks that overlapped binding sites of the corresponding TF.

### – Classification of TF-Dependent Regulatory Outcomes

To classify TF-dependent promoter regulation, we integrated differential ATAC-seq and RNA-seq responses following 4 h degradation of BANP, GABPA, or ZFP143. TF-dependent ATAC targets were defined as promoter-overlapping ATAC-seq peaks significantly altered by TF depletion (DESeq2, adjusted P < 0.05) and intersecting a verified TF ChIP-seq peak. TF-dependent RNA targets were defined as genes significantly up- or down-regulated upon depletion (DESeq2, adjusted P < 0.01). For each TF, genes were assigned to one of four mutually exclusive classes based on the combination of accessibility and expression changes at their promoters: ATAC+RNA, ATAC-only, RNA-only, or No effect. These classifications were used for all downstream analyses of TF regulatory logic and co-regulation.

### – TOBIAS

Footprinting and Differential Binding TOBIAS (v0.17.1, https://github.com/loosolab/TOBIAS) was used to correct ATAC–seq signal for Tn5 bias, compute footprint scores, and quantify differential TF binding. BAM files generated from the ATAC-seq pipeline (as described above) were processed using ATACorrect (bias correction using the reference genome and unified TF peak set) and FootprintScores (generation of per-base footprint bigWigs). Differential TF binding was assessed with BINDetect, using JASPAR motif collections and the corrected footprint signals as input with pairwise comparisons between UT and 4 h samples. BINDetect reported motif occurrences, footprint strength, and differential binding statistics for each TF.

### – Motif Variant Analysis

To assess whether promoter classes differ in motif quality, we first scanned the unified set of merged TF-bound peaks using FIMO with the corresponding TF PWMs reported in JASPAR. Motif occurrences overlapping promoter regions were assigned to their regulatory class (e.g., TF1-specific, co-regulated, TF2-specific). For each class, the underlying DNA sequences were collected and used to generate class-specific position frequency matrices (PFMs), which were then converted to position weight matrices (PWMs) using the universalmotif package (https://github.com/bjmt/universalmotif). Motif strength was obtained by computing the average FIMO score across all motif instances within that category. This approach enabled direct evaluation of whether functional promoter categories are associated with systematic differences in motif strength.

### – Human Population Constraint Analysis of TF Motifs

To assess population-level constraint on TF motifs in human promoters, we analyzed genome-wide variants from gnomAD v4.0. Chromosome-resolved VCF files for all autosomes and sex chromosomes were downloaded from the gnomAD public Google Cloud repository. Variants overlapping promoter regions were extracted and assigned to essential or non-essential gene groups based on the classification described above. These promoter variants were then intersected with TF motif coordinates for BANP, GABPA, MYC, YY1, and ZFP143, defined around human TSSs, yielding motif-annotated variant tables that included genomic position, alleles, gene identity, motif name, and allele frequency when available. For each TF and gene essentiality class, we quantified the total genomic span occupied by that motif (summed motif lengths across essential or non-essential promoters) and normalized the number of overlapping variants by this value to compute a variant density (variants per kilobase of motif sequence). Population constraint was evaluated by comparing variant densities between essential and non-essential promoters and by performing Fisher’s exact tests to compute odds ratios and associated P values.

### – PromoterAI variant impact analysis

We obtained pre-computed promoterAI scores from the original authors, providing predicted transcriptional effects for all possible single-nucleotide variants within ±500 bp of every human TSS. Variant entries were filtered to retain only promoters of genes of interest, and genomic coordinates were converted to BED format. Filtered promoterAI variants were intersected with TF motif coordinates (BANP, GABPA, ZFP143, YY1, MYC) to annotate whether each variant overlapped a predicted TF-binding site. This produced a unified table listing, for each variant, its promoterAI score, gene identity, and TF motif overlap. Genes were assigned to regulatory classes (ATAC-only, ATAC+RNA, RNA-only, non-responsive) based on classifications described above, and their variants were categorized accordingly. For each class and TF, promoterAI scores were summarized and compared using non-parametric tests (Wilcoxon or Kruskal–Wallis), and distributions were visualized using violin and boxplots.

#### Single promoter plots

For a given gene, we first selected a single representative TSS in a strand-aware manner (most 5′ genomic position for + strand, most 3′ for − strand) and recalculated all variant positions relative to this TSS. We then restricted the analysis to variants within −200 to +200 bp of the TSS. At each genomic position, we identified the reference base and averaged the promoterAI scores across all possible alternative alleles at that base, yielding one mean promoterAI value per position. These values were assembled into a 4×N matrix (A/C/G/T × position), with only the reference base at each position assigned the corresponding mean promoterAI score. The matrix was normalized by the maximum absolute value and sign-inverted so that larger positive values represent positions where variants are predicted to have stronger negative impact on promoter activity. This matrix was visualized as a signed DNA sequence logo (ggseqlogo), aligned to a line plot of the mean promoterAI score as a function of distance from the TSS, providing a position-resolved view of promoter sensitivity to sequence perturbation.

## References

1. Hart, T. et al. High-Resolution CRISPR Screens Reveal Fitness Genes and Genotype-Specific Cancer Liabilities. Cell 163, 1515–26 (2015).

2. Wang, T. et al. Identification and characterization of essential genes in the human genome. Science 350, 1096–101 (2015).

3. Jeffries, A. M. et al. Single-cell transcriptomic and genomic changes in the ageing human brain. Nature 646, 657–666 (2025).

4. Dhanasekaran, R. et al. The MYC oncogene - the grand orchestrator of cancer growth and immune evasion. Nat. Rev. Clin. Oncol. 19, 23–36 (2022).

5. López-Otín, C., Blasco, M. A., Partridge, L., Serrano, M. & Kroemer, G. Hallmarks of aging: An expanding universe. Cell 186, 243–278 (2023).

6. Deaton, A. M. & Bird, A. CpG islands and the regulation of transcription. Genes Dev. 25, 1010–22 (2011).

7. Blackledge, N. P. & Klose, R. CpG island chromatin: a platform for gene regulation. Epigenetics 6, 147–52 (2011).

8. Lambert, S. A. et al. The Human Transcription Factors. Cell 172, 650–665 (2018).

9. Todeschini, A.-L., Georges, A. & Veitia, R. A. Transcription factors: specific DNA binding and specific gene regulation. Trends Genet. 30, 211–9 (2014).

10. Slattery, M. et al. Absence of a simple code: how transcription factors read the genome. Trends Biochem. Sci. 39, 381–99 (2014).

11. Zhu, F. et al. The interaction landscape between transcription factors and the nucleosome. Nature 562, 76–81 (2018).

12. Grand, R. S. et al. Genome access is transcription factor-specific and defined by nucleosome position. Mol. Cell 84, 3455–3468.e6 (2024).

13. Zaret, K. S. & Carroll, J. S. Pioneer transcription factors: establishing competence for gene expression. Genes Dev. 25, 2227–41 (2011).

14. Isbel, L., Grand, R. S. & Schübeler, D. Generating specificity in genome regulation through transcription factor sensitivity to chromatin. Nat. Rev. Genet. 23, 728–740 (2022).

15. Burger, L., Gil, N. & Schübeler, D. A novel deep learning-based framework reveals a continuum of chromatin sensitivities across transcription factors. Preprint at 10.1101/2025.08.11.669605 (2025).

16. Spitz, F. & Furlong, E. E. M. Transcription factors: from enhancer binding to developmental control. Nat. Rev. Genet. 13, 613–26 (2012).

17. Kim, S. et al. DNA-guided transcription factor cooperativity shapes face and limb mesenchyme. Cell 187, 692–711.e26 (2024).

18. Morgunova, E. & Taipale, J. Structural perspective of cooperative transcription factor binding. Curr. Opin. Struct. Biol. 47, 1–8 (2017).

19. Sönmezer, C. et al. Molecular Co-occupancy Identifies Transcription Factor Binding Cooperativity In Vivo. Mol. Cell 81, 255–267.e6 (2021).

20. Jolma, A. et al. DNA-dependent formation of transcription factor pairs alters their binding specificity. Nature 527, 384–8 (2015).

21. Soufi, A. et al. Pioneer transcription factors target partial DNA motifs on nucleosomes to initiate reprogramming. Cell 161, 555–568 (2015).

22. Michael, A. K. et al. Mechanisms of OCT4-SOX2 motif readout on nucleosomes. Science 368, 1460–1465 (2020).

23. Kim, J. & DeBerardinis, R. J. Mechanisms and Implications of Metabolic Heterogeneity in Cancer. Cell Metab. 30, 434–446 (2019).

24. Gurumayum, S. et al. OGEE v3: Online GEne Essentiality database with increased coverage of organisms and human cell lines. Nucleic Acids Res. 49, D998–D1003 (2021).

25. Arafeh, R., Shibue, T., Dempster, J. M., Hahn, W. C. & Vazquez, F. The present and future of the Cancer Dependency Map. Nat. Rev. Cancer 25, 59–73 (2025).

26. Hammal, F., de Langen, P., Bergon, A., Lopez, F. & Ballester, B. ReMap 2022: a database of Human, Mouse, Drosophila and Arabidopsis regulatory regions from an integrative analysis of DNA-binding sequencing experiments. Nucleic Acids Res. 50, D316–D325 (2022).

27. Grand, R. S. et al. BANP opens chromatin and activates CpG-island-regulated genes. Nature 596, 133–137 (2021).

28. Fan, K., Moore, J. E., Zhang, X.-O. & Weng, Z. Genetic and epigenetic features of promoters with ubiquitous chromatin accessibility support ubiquitous transcription of cell-essential genes. Nucleic Acids Res. 49, 5705–5725 (2021).

29. Zhou, C., Wang, M., Zhang, C. & Zhang, Y. The transcription factor GABPA is a master regulator of naive pluripotency. Nat. Cell Biol. 27, 48–58 (2025).

30. Liu, B. B. et al. Dissecting regulatory syntax in human development with scalable multiomics and deep learning. Preprint at 10.1101/2025.04.30.651381 (2025).

31. Jaganathan, K. et al. Predicting expression-altering promoter mutations with deep learning. Science 389, eads7373 (2025).

32. Rauluseviciute, I. et al. JASPAR 2024: 20th anniversary of the open-access database of transcription factor binding profiles. Nucleic Acids Res. 52, D174–D182 (2024).

33. Jinek, M. et al. A programmable dual-RNA-guided DNA endonuclease in adaptive bacterial immunity. Science 337, 816–21 (2012).

34. Weintraub, A. S. et al. YY1 Is a Structural Regulator of Enhancer-Promoter Loops. Cell 171, 1573–1588.e28 (2017).

35. Narducci, D. N. & Hansen, A. S. Putative looping factor ZNF143/ZFP143 is an essential transcriptional regulator with no looping function. Mol. Cell 85, 9–23.e9 (2025).

36. Zhou, C., Wang, M., Zhang, C. & Zhang, Y. The transcription factor GABPA is a master regulator of naive pluripotency. Nat. Cell Biol. 27, 48–58 (2025).

37. Fernandez, P. C. et al. Genomic targets of the human c-Myc protein. Genes Dev. 17, 1115–29 (2003).

38. Duttke, S. H. et al. Position-dependent function of human sequence-specific transcription factors. Nature 631, 891–898 (2024).

39. Sahu, B. et al. Sequence determinants of human gene regulatory elements. Nat. Genet. 54, 283–294 (2022).

40. Dudnyk, K., Cai, D., Shi, C., Xu, J. & Zhou, J. Sequence basis of transcription initiation in the human genome. Science 384, eadj0116 (2024).

41. Nabet, B. et al. The dTAG system for immediate and target-specific protein degradation. Nat. Chem. Biol. 14, 431–441 (2018).

42. Gaidatzis, D., Burger, L., Florescu, M. & Stadler, M. B. Analysis of intronic and exonic reads in RNA-seq data characterizes transcriptional and post-transcriptional regulation. Nat. Biotechnol. 33, 722–9 (2015).

43. Mahendrawada, L., Warfield, L., Donczew, R. & Hahn, S. Low overlap of transcription factor DNA binding and regulatory targets. Nature 642, 796–804 (2025).

44. Isbel, L. et al. Readout of histone methylation by Trim24 locally restricts chromatin opening by p53. Nat. Struct. Mol. Biol. 30, 948–957 (2023).

45. Buenrostro, J. D., Wu, B., Chang, H. Y. & Greenleaf, W. J. ATAC-seq: A Method for Assaying Chromatin Accessibility Genome-Wide. Curr. Protoc. Mol. Biol. 109, 21.29.1–21.29.9 (2015).

46. Bond, A. G. et al. Development of BromoTag: A ‘Bump-and-Hole’-PROTAC System to Induce Potent, Rapid, and Selective Degradation of Tagged Target Proteins. J. Med. Chem. 64, 15477–15502 (2021).

47. Bentsen, M. et al. ATAC-seq footprinting unravels kinetics of transcription factor binding during zygotic genome activation. Nat. Commun. 11, 4267 (2020).

48. Grand, R. S. et al. BANP opens chromatin and activates CpG-island-regulated genes. Nature 596, 133–137 (2021).

49. Love, M. I., Huber, W. & Anders, S. Moderated estimation of fold change and dispersion for RNA-seq data with DESeq2. Genome Biol. 15, 550 (2014).

50. Lienert, F. et al. Identification of genetic elements that autonomously determine DNA methylation states. Nat. Genet. 43, 1091–7 (2011).

51. Chen, S. et al. A genomic mutational constraint map using variation in 76,156 human genomes. Nature 625, 92–100 (2024).

52. Patange, S. et al. MYC amplifies gene expression through global changes in transcription factor dynamics. Cell Rep. 38, 110292 (2022).

53. Dratwa, M., Wysoczańska, B., Łacina, P., Kubik, T. & Bogunia-Kubik, K. TERT-Regulation and Roles in Cancer Formation. Front. Immunol. 11, 589929 (2020).

54. Heidenreich, B., Rachakonda, P. S., Hemminki, K. & Kumar, R. TERT promoter mutations in cancer development. Curr. Opin. Genet. Dev. 24, 30–7 (2014).

55. Wu, K. J. et al. Direct activation of TERT transcription by c-MYC. Nat. Genet. 21, 220–4 (1999).

56. Andersson, R. & Sandelin, A. Determinants of enhancer and promoter activities of regulatory elements. Nat. Rev. Genet. 21, 71–87 (2020).

57. Deaton, A. M. & Bird, A. CpG islands and the regulation of transcription. Genes Dev. 25, 1010–1022 (2011).

58. Babu, S., Takeuchi, Y. & Masai, I. Banp regulates DNA damage response and chromosome segregation during the cell cycle in zebrafish retina. Elife 11, (2022).

59. Xiong, L. et al. Oct4 differentially regulates chromatin opening and enhancer transcription in pluripotent stem cells. Elife 11, (2022).

60. Maresca, M. et al. Pioneer activity distinguishes activating from non-activating SOX2 binding sites. EMBO J. 42, e113150 (2023).

61. Domcke, S. et al. Competition between DNA methylation and transcription factors determines binding of NRF1. Nature 528, 575–9 (2015).

62. Kim, J. Do, Faulk, C. & Kim, J. Retroposition and evolution of the DNA-binding motifs of YY1, YY2 and REX1. Nucleic Acids Res. 35, 3442–52 (2007).

63. Barral, A. & Zaret, K. S. Pioneer factors: roles and their regulation in development. Trends Genet. 40, 134–148 (2024).

64. Ouyang, Z. et al. The developmental and evolutionary characteristics of transcription factor binding site clustered regions based on an explainable machine learning model. Nucleic Acids Res. 52, 7610–7626 (2024).

65. Domcke, S. et al. Competition between DNA methylation and transcription factors determines binding of NRF1. Nature 528, 575–9 (2015).

66. Durdu, S. et al. Chromatin-dependent motif syntax defines differentiation trajectories. Mol. Cell 85, 2900–2918.e16 (2025).

67. Lupo, O. et al. The architecture of binding cooperativity between densely bound transcription factors. Cell Syst. 14, 732–745.e5 (2023).

68. Illingworth, R. S. & Bird, A. P. CpG islands--’a rough guide’. FEBS Lett. 583, 1713–20 (2009).

69. Deaton, A. M. & Bird, A. CpG islands and the regulation of transcription. Genes Dev. 25, 1010–22 (2011).

70. Sankar, A. et al. Histone editing elucidates the functional roles of H3K27 methylation and acetylation in mammals. Nat. Genet. 54, 754–760 (2022).

71. Hughes, A. L. et al. A CpG island-encoded mechanism protects genes from premature transcription termination. Nat. Commun. 14, 726 (2023).

72. Policarpi, C., Munafò, M., Tsagkris, S., Carlini, V. & Hackett, J. A. Systematic epigenome editing captures the context-dependent instructive function of chromatin modifications. Nat. Genet. 56, 1168–1180 (2024).

73. Fukai, Y. T. et al. Gene-scale in vitro reconstitution reveals histone acetylation directly controls chromatin architecture. Sci. Adv. 11, eadx9282 (2025).

74. Weilert, M. et al. Widespread low-affinity motifs enhance chromatin accessibility and regulatory potential in mESCs. Preprint at 10.1101/2025.11.18.685822 (2025).

75. Sasse, A. et al. Benchmarking of deep neural networks for predicting personal gene expression from DNA sequence highlights shortcomings. Nat. Genet. 55, 2060–2064 (2023).

76. Barbadilla-Martínez, L., Klaassen, N., van Steensel, B. & de Ridder, J. Predicting gene expression from DNA sequence using deep learning models. Nat. Rev. Genet. 26, 666–680 (2025).

77. Weber, M. et al. Distribution, silencing potential and evolutionary impact of promoter DNA methylation in the human genome. Nat. Genet. 39, 457–66 (2007).

78. Gao, T. & Qian, J. EnhancerAtlas 2.0: an updated resource with enhancer annotation in 586 tissue/cell types across nine species. Nucleic Acids Res. 48, D58–D64 (2020).

79. Zhou, Q. et al. A mouse tissue transcription factor atlas. Nat. Commun. 8, 15089 (2017).

