## Supplemental Figures for "Essential genes are dominantly activated by single transcription factors"

### Extended data

#### Extended data Fig.1

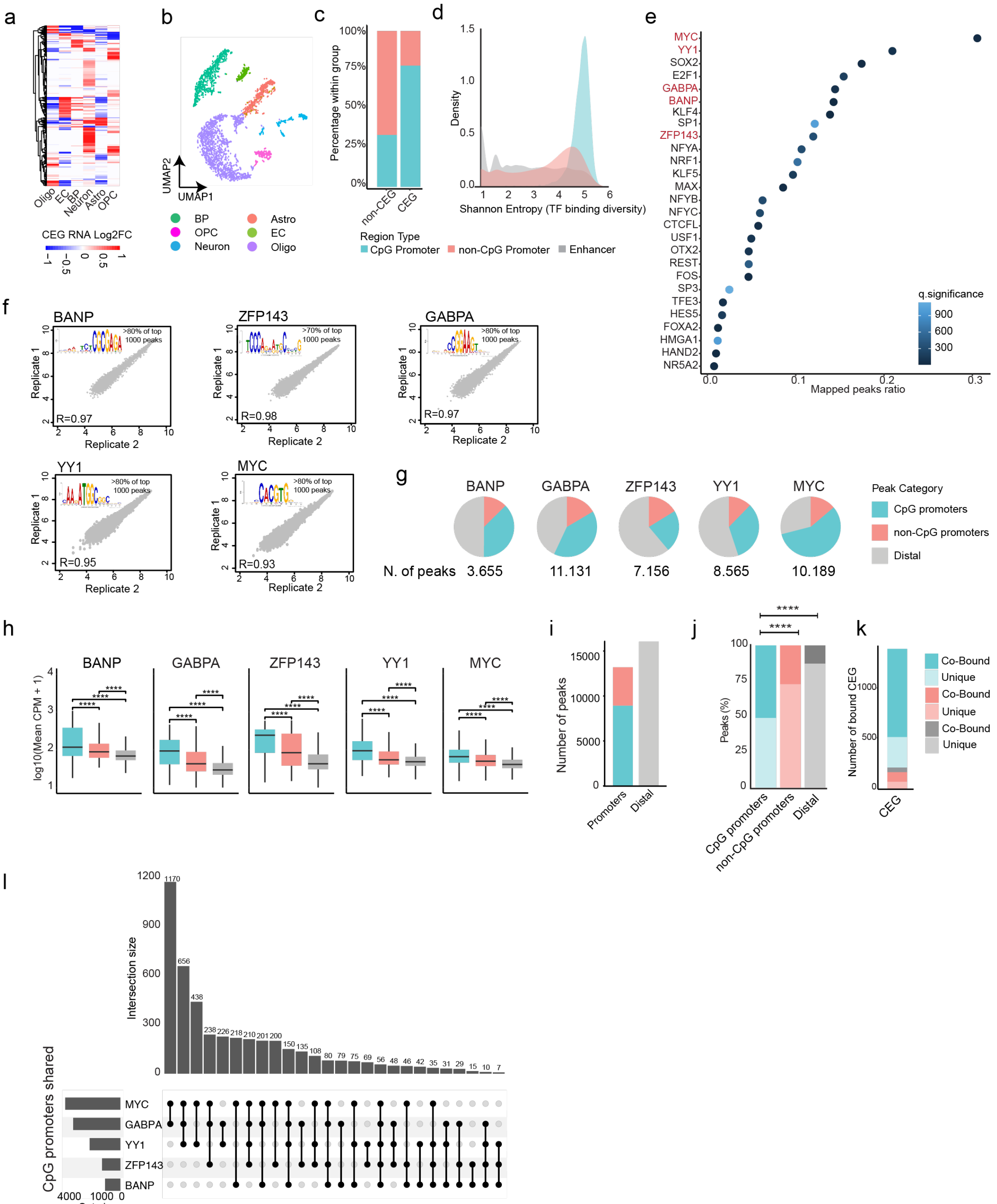


**Extended data Fig.1. TFs co-bind extensively around the TSS of CGI-promoter-regulated genes.** a) Differential expression of 408 CEGs across the different cell types in the brain. b) Total gene expression profiles enable separation of brain cell types. Analysis was performed using Tabula Muris single-cell RNA-seq data. c) Distribution of promoter class (CpG / non-CpG) in CEG or non-CEG. d) TF binding diversity across distinct classes of cis-regulatory elements (CREs), measured using Shannon entropy. e) The fraction of TF binding that overlaps with CEG promoters and have a motif using ReMap data. f) Scatter plots showing

ChIP–seq replicate correlations. Pearson correlation coefficients exceeded 0.9 for all TFs. Motif enrichment analysis using MEME suite on the top 1,000 peaks identified canonical TF motifs. g) Pie charts showing the distribution of TF binding sites across CpG island promoters (CGI), non-CpG promoters, and distal elements. h) Box plots of TF binding intensity within their respective peaks, stratified by CRE category. I) and j) Distribution of a merged set of ~29,000 TF peaks across CRE categories. k) Stacked bar plots showing the proportion of binding sites where each TF binds alone (1 TF) or co-binds with at least one other TF (≥2 TFs) across CRE categories. Significance indicates a p-value =< 0.001. K) UpSet plot illustrating specific TF co-binding combinations. I) The distribution of CEGs across the different CREs and co-binding classes.

#### Extended data Fig.2

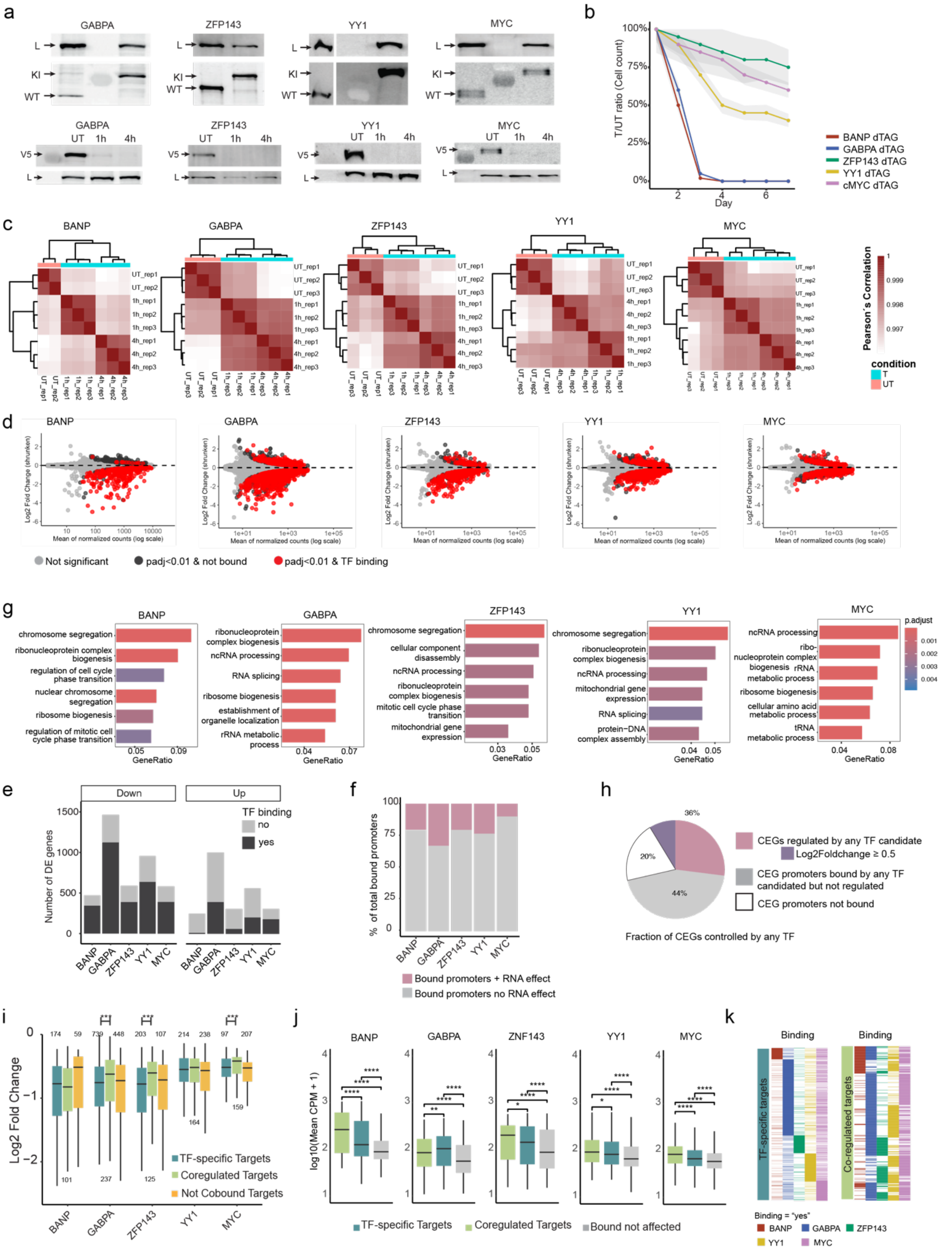


**Extended data Fig.2. Co-bound genes are predominantly controlled by a single TF**. a) Western blot of homozygous knockin clones showing depletion following one hour of degradation upon dTAG13 treatment. b) Cell viability following extended TF degradation. Grey shadow represents +/-SD. c) Heatmap showing Pearson correlation between RNA-seq replicates, confirming high reproducibility. d) MA plots of gene expression changes ($\log_2$ fold change vs. base mean of intronic counts). Colored dots indicate genes whose promoters are bound by a given TF and are significantly affected by its depletion. e) Stacked bar plots showing the number of up- and downregulated genes per TF upon degradation, classified in bound by the respective TF at the promoter of the same gene or not bound. f) Stacked bar plots showing the distribution of TF binding sites between genes that are differentially expressed upon TF removal and those that are not. g) Gene Ontology (GO) enrichment analysis of primary target genes regulated by each TF. h) Pie chart showing the proportion of CEGs that are: (1) bound and regulated by at least one candidate TF, (2) bound but not regulated, or (3) not bound by any candidate TF. i) Box plots showing fold changes ($log_2$) in gene expression following TF removal, grouped by regulatory categories. j) Box plots showing TF binding intensity (counts per million, CPM) at co-bound promoters of genes that are co-regulated, uniquely regulated by a single TF, or not transcriptionally affected. k) Heatmap showing the binding of each of the TFs at the individual and co-regulated gene promoters, ranked as in Figure 2e and f. Colours represent the different TFs. * = pvalue<0.01, *** = pvalue<0.0001.

#### Extended data Fig.3

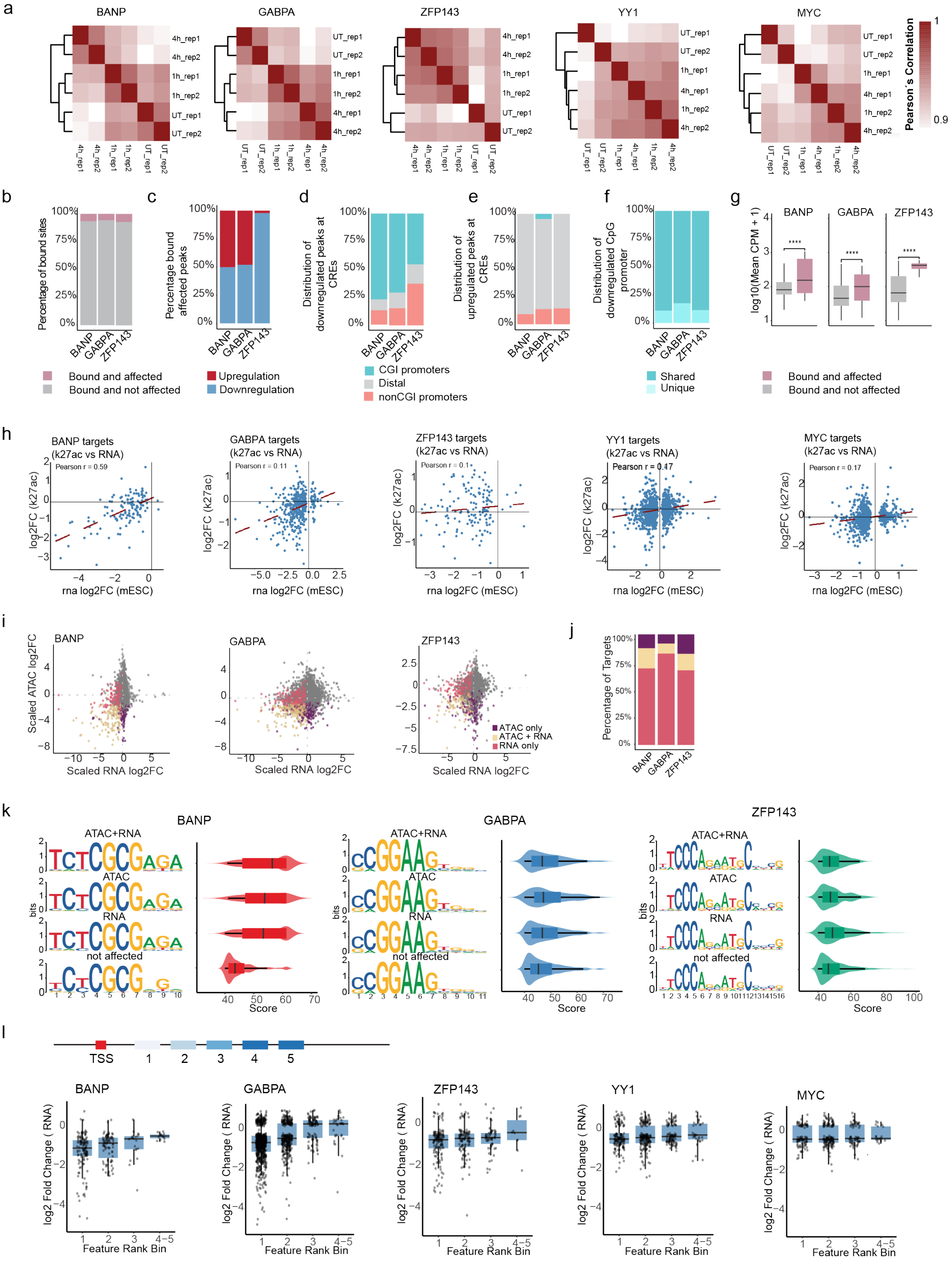


**Extended data Fig.3. Chromatin organization and modification at CGI promoters**. a) Heatmap showing Pearson correlation between ATAC-seq replicates, demonstrating high reproducibility. b) Stacked bar plots showing the distribution of TF binding sites across peaks that are differentially accessible upon TF depletion and those that remain unchanged. c) Stacked bar plots showing the number of peaks that gain or lose accessibility following TF degradation. d) Distribution of peaks that lose accessibility upon TF removal, stratified by cis-regulatory element (CRE) category. e) Distribution of peaks that gain accessibility upon TF removal, stratified by CRE category. f) Distribution of peaks that lose accessibility at CpG island (CGI) promoters, classified as uniquely or jointly bound by TFs. g) Box plots comparing TF binding intensity (counts per million, CPM) at peaks with or without accessibility changes upon TF loss. h) Scatter plot showing correlation between $\log_2$ fold changes in H3K27ac (ChIP–seq) and gene expression (RNA-seq) upon individual TF depletion. i) Scatter plot showing correlation between changes in promoter accessibility (ATAC-seq $\log_2$ fold change) and gene expression (RNA-seq) upon TF removal. Promoters are classified into three categories: ATAC-only, RNA-only, or ATAC+RNA responsive. j) Stacked bar plots showing the distribution of TF-bound promoters across the categories defined in J. Promoters in the ATAC+RNA group are further subdivided into genes that are co-regulated by multiple TFs or specifically regulated by a single TF. k) Violin plots showing the distribution of relative motif scores for motif variants enriched at functional (ATAC+RNA, RNA-only, ATAC-only) versus non-functional TF binding sites. l) Schematic illustrating the classification of TF binding positions relative to the TSS and co-bound factors (top). Promoter-associated target genes were grouped according to the relative position of TF binding, and boxplots display the corresponding $\log_2$ fold change in RNA expression following depletion of each TF (bottom). *** = $p<0.0001$.

**Extended data Fig.4**

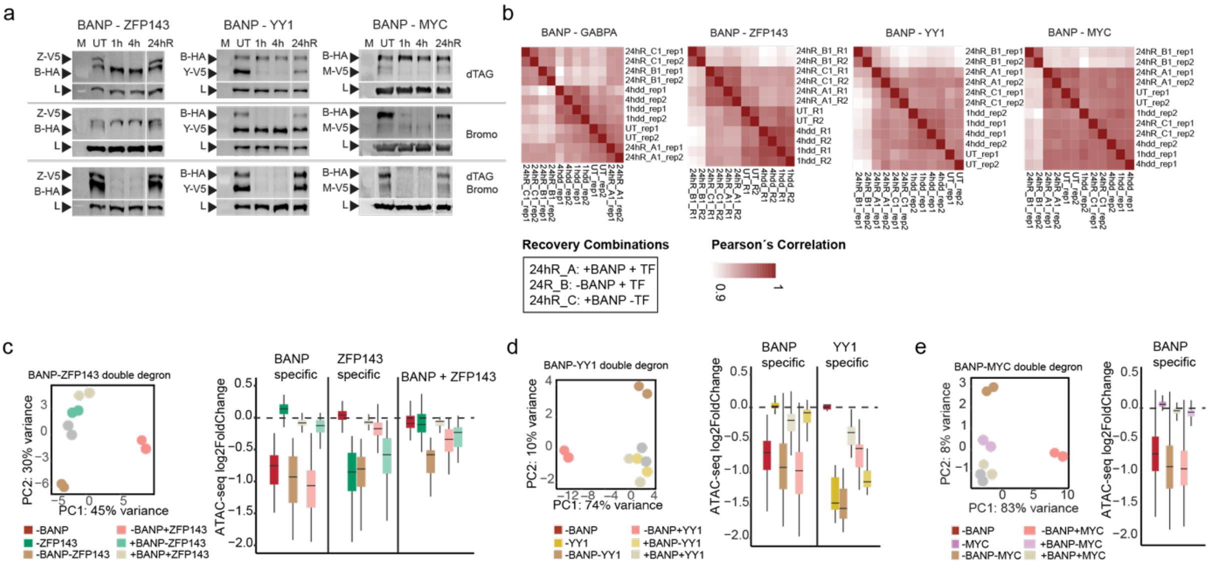


**Extended data Fig.4. Individual TFs drive chromatin opening at specific genes.** a) Western blot analysis of BANP and TF protein levels after 1 h, 2 h, and 4 h of dTAG13 and/or AGB1 treatment, demonstrating efficient protein depletion and recovery. VCL was used as a loading control. b) Heatmap showing Pearson correlation between ATAC-seq replicates, demonstrating high reproducibility. c) PCA of ATAC-seq data showing accessibility divergence after both TF degradation, recovery of only ZFP143, recovery of only BANP, and recovery of both TFs, relative to untreated (UT). Right: boxplots displaying $\log_2$ fold-changes in accessibility (ATAC-seq) for peaks classified as BANP-specific, ZFP143-specific, or BANP+ ZFP143-dependent following individual and combinatorial degradation and recovery conditions. Centre line, median; box limits, upper and lower quartiles; whiskers, minimum and maximum. d) PCA of ATAC-seq data showing accessibility divergence after both TF degradation, recovery of only YY1, recovery of only BANP, and recovery of both TFs, relative to untreated (UT). Corresponding boxplots displaying $\log_2$ fold-changes in accessibility (ATAC-seq) for peaks classified as BANP-specific, YY1-specific, following individual and combinatorial degradation and recovery conditions (statistics as in c). e) PCA of ATAC-seq data showing accessibility divergence after both TF degradation, recovery of only MYC, recovery of only BANP, and recovery of both TFs, relative to untreated (UT). Corresponding boxplots displaying $\log_2$ fold-changes in accessibility (ATAC-seq) for peaks classified as BANP-specific following individual and combinatorial degradation and recovery conditions. (statistics as in c).

**Extended data Fig.5**

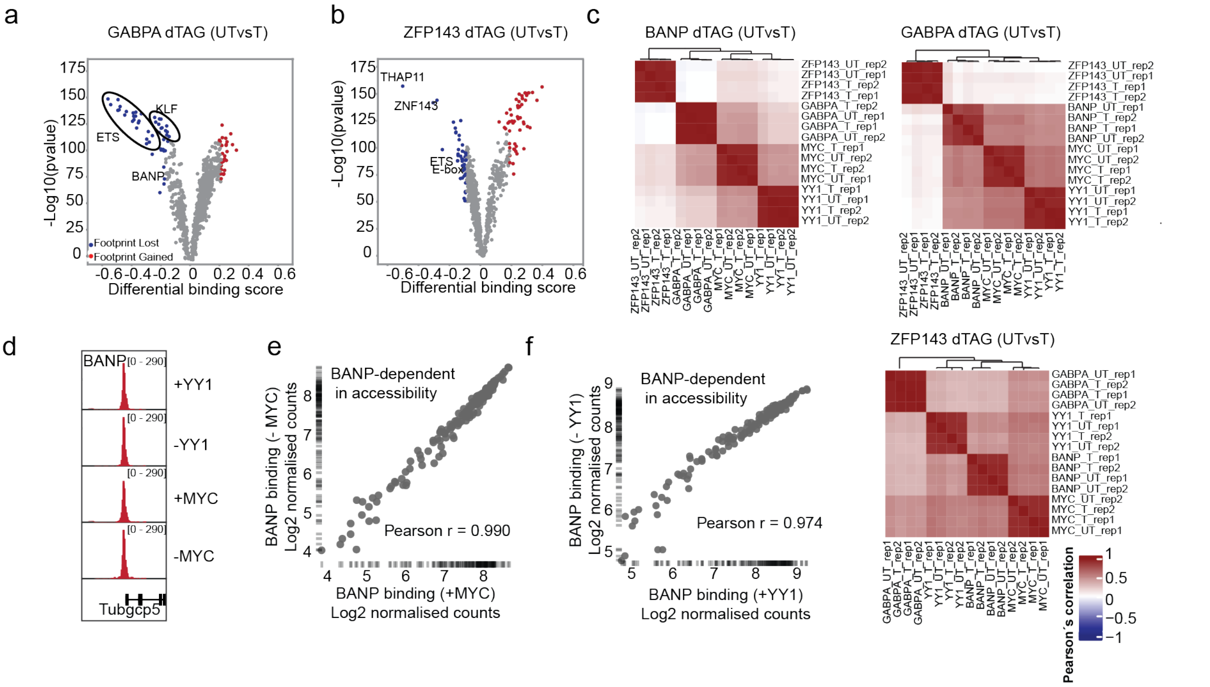


**Extended data Fig.5: Chromatin organization impacts the co-binding of TFs.** a) Volcano plot displaying differential TF motif occupancy upon GABPA degradation, as assessed by TOBIAS footprinting. Binding scores are plotted for individual motifs, highlighting significant changes in accessibility following GABPA loss. b) Volcano plot displaying differential TF motif occupancy upon ZFP143 degradation, as assessed by TOBIAS footprinting. Binding scores are plotted for individual motifs, highlighting significant changes in accessibility following ZFP143 loss. c) Heatmap showing Pearson correlation between ChIP-seq replicates in two conditions: before TF degradation (UT) and after 1h of TF degradation (T). d) Single locus example of BANP binding at the Tubgcp5 promoter, before and after removal of GABPA, YY1, or MYC. e) Scatterplot showing correlation of BANP ChIP raw counts performed before and after removing cMYC. Person's correlation is indicated. f) Same as e) but before and after removing YY1.

#### Extended data Fig.6

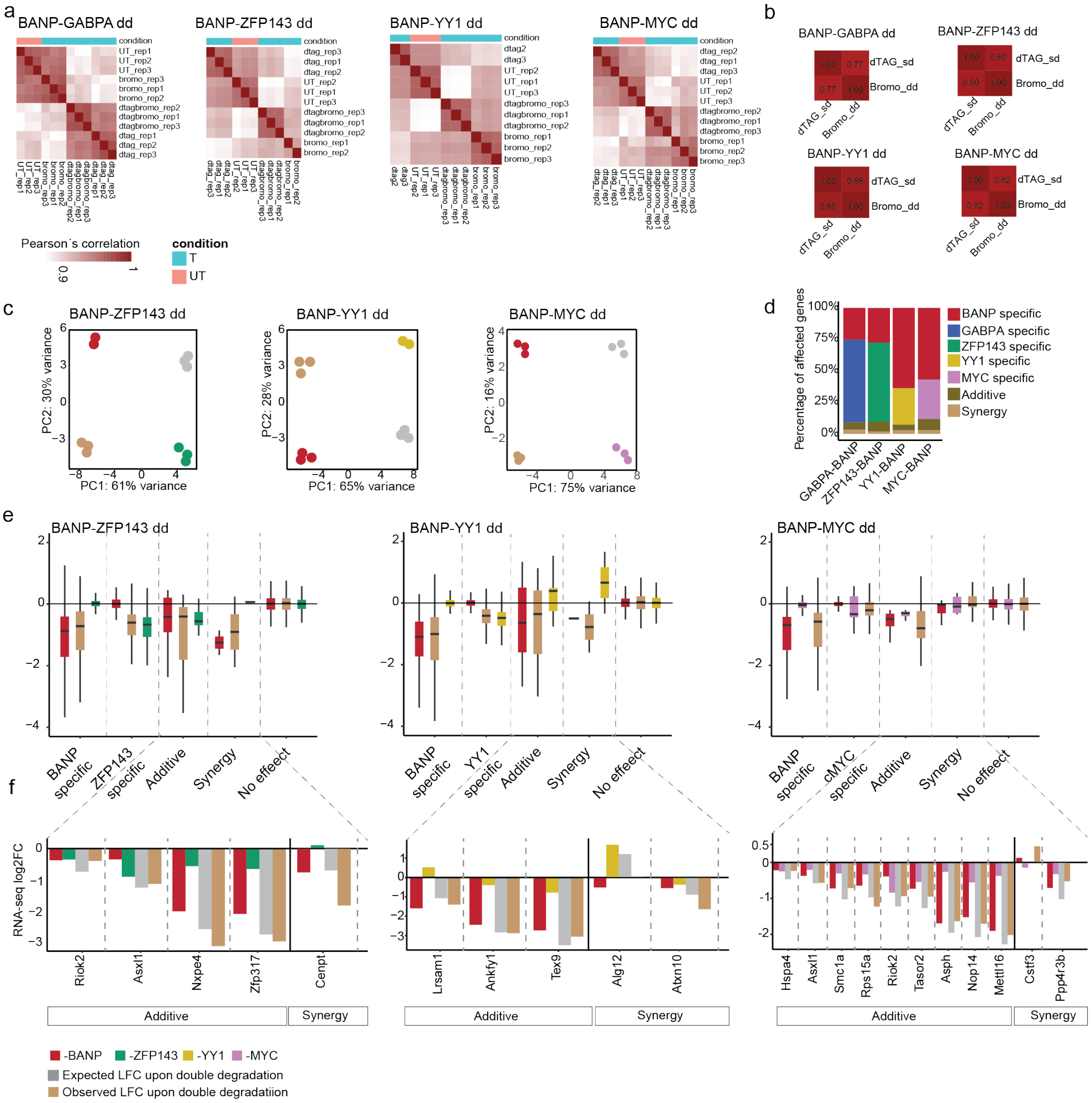


**Extended data Fig.6. Co-binding of TFs can tune gene activity at a small set of loci**. a) Heatmap showing Pearson correlation between RNA-seq replicates in four conditions: before TF degradation (UT), after 4h of BANP degradation (bromo), after 4h of TF degradation (dtag), after double degradation (dtag_bromo). b) Heatmap showing Pearson correlation between Log2FoldChange induced by BANP degradation with dTAG or with Bromo across the different double degron lines, restricted to all primary BANP target genes (defined in the single degron experiments). c) PCA of RNA-seq data showing transcriptional divergence after the degradation of individual or both TFs, relative to untreated (UT). d) The fraction of genes significantly impacted by the individual or combinatorial removal of different TF pairs. e) Box plots showing the fold-change (log2) of the different responsive gene classes. f) Bar plots showing gene expression changes (log2 fold change) upon single or double TF degradation for all genes detected by interaction contrast analysis. Genes are grouped into two categories based on interaction contrast: additive-co-regulated and synergistic.

#### Extended data Fig.7

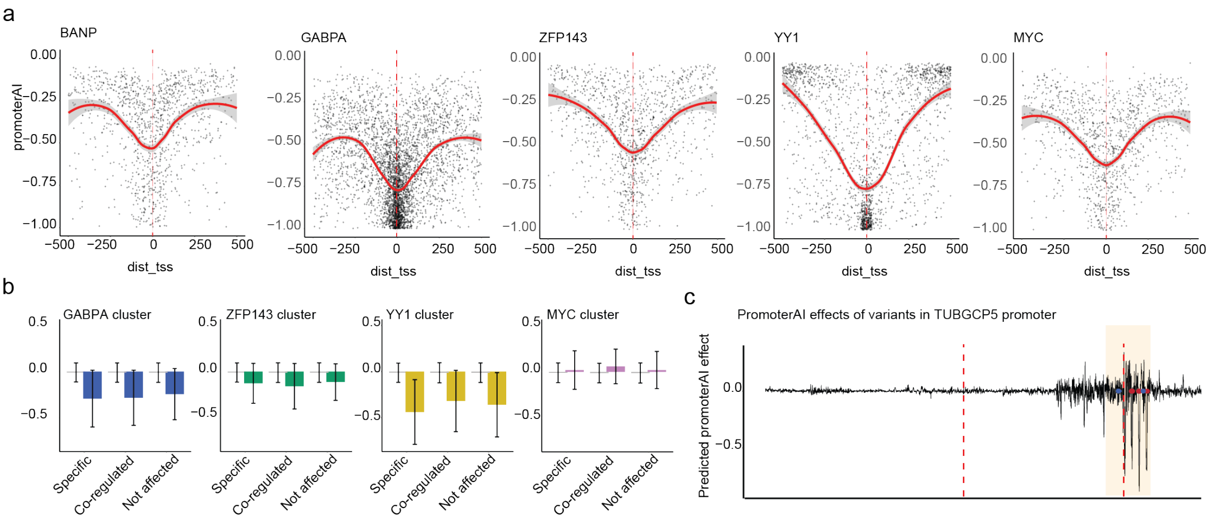


**Extended data Fig.7. Conservation of regulatory dominance in human cells.** a) Mean predicted variant effects (promoterAI scores) at specific TF motifs grouped by distance to the TSS. Shaded areas indicate the standard error of the mean. b) Average promoterAI scores at core motifs within TF clusters specific, co-regulated, not affected, for each TF (GABPA, ZFP143, YY1, and MYC). c) Mean of predicted promoterAI score for the TUBGCP5 promoter. The plot shows a 1 kb region centered on the transcription start site (dashed red line), with a highlighted window marking the BANP motif region where variant effects are concentrated.
